# A genome-wide CRISPR screen reveals a role for the BRD9-containing non-canonical BAF complex in regulatory T cells

**DOI:** 10.1101/2020.02.26.964981

**Authors:** Chin-San Loo, Jovylyn Gatchalian, Yuqiong Liang, Mathias Leblanc, Mingjun Xie, Josephine Ho, Bhargav Venkatraghavan, Diana C. Hargreaves, Ye Zheng

## Abstract

Regulatory T cells (Tregs) play a pivotal role in suppressing auto-reactive T cells and maintaining immune homeostasis. Treg development and function are dependent on the transcription factor Foxp3. Here we performed a genome-wide CRISPR/Cas9 knockout screen to identify the regulators of Foxp3 in mouse primary Tregs. The results showed that Foxp3 regulators are highly enriched in genes encoding SWI/SNF and SAGA complex subunits. Among the three SWI/SNF-related complexes, the non-canonical or ncBAF (also called GBAF or BRD9-containing BAF) complex promoted the expression of Foxp3, whereas the PBAF complex repressed its expression. Gene ablation of BRD9 led to compromised Treg function in inflammatory disease and tumor immunity. Functional genomics revealed that BRD9 is required for Foxp3 binding and expression of a subset of Foxp3 target genes. Thus, we provide an unbiased analysis of genes and networks regulating Foxp3, and reveal ncBAF complex as a novel target that could be exploited to manipulate Treg function.

## Introduction

Regulatory T cells (Tregs) play a crucial role in maintaining immune system homeostasis by suppressing over-reactive immune responses(Josefowicz et al., 2012; Sakaguchi et al., 2008). Defects in Tregs lead to autoimmune disorders and immunopathology, while certain tumors are enriched with Tregs that suppress anti-tumor immune responses(Tanaka and Sakaguchi, 2017). Foxp3, a member of the Forkhead transcription factor family, is a critical regulator that orchestrates the molecular processes involved in Treg differentiation and function(Zheng and Rudensky, 2007). Therefore, understanding the regulation of Foxp3 expression could reveal novel therapeutic targets to potentially change Treg numbers or alter their function. It has been established that the T cell receptor (TCR) and IL-2 signaling pathways play critical roles in Foxp3 induction(Chinen et al., 2016; Lee et al., 2012). TGF-β signaling is also essential for Foxp3 induction in periphery-derived Tregs and in vitro induced Tregs, although its role in thymus-derived Treg development is still under debate (Chen et al., 2003; Liu et al., 2008; Ouyang et al., 2010). Accordingly, a number of downstream transcription factors have been identified that regulate Foxp3 induction in vitro or in vivo, including STAT5a/b, CBF-β/RUNX1/3, NFAT1, SMAD3/4, cRel, and CREB (Burchill et al., 2007; Kim and Leonard, 2007; Kitoh et al., 2009; Long et al., 2009; Rudra et al., 2009; Tone et al., 2008; Yang et al., 2008). Compared to the large number of studies focused on the mechanism of Foxp3 induction, relatively less is known about the factors that maintain Foxp3 expression in mature Treg cells. An intronic enhancer in *Foxp3* named CNS2 (conserved non-coding sequence 2), also known as TSDR (Treg-specific demethylated region), is a key cis-regulatory element required for stable Foxp3 expression(Polansky et al., 2008; Zheng et al., 2010). CNS2 is heavily methylated in naive and activated conventional T cells by DNA methyl-transferase 1 (DNMT1), and deletion of Dnmt1 leads to aberrant expression of Foxp3 in conventional T cells(Josefowicz et al., 2009). Once Foxp3 expression is induced during Treg development, the CNS2 region is rapidly demethylated, opening it up for the binding of transcription factors(Polansky et al., 2008). Foxp3 can bind to CNS2, as well as an additional upstream enhancer named CNS0(Kitagawa et al., 2017), and stabilize its own expression in a positive feedback loop(Feng et al., 2014; Li et al., 2014b).

Post-translational modifications (PTM) of the Foxp3 protein, including phosphorylation, acetylation, and ubiquitination, are also a crucial part of the regulatory circuit that controls Foxp3 function and stability (van Loosdregt and Coffer, 2014). Among the regulators of Foxp3 PTMs, a pair of enzymes, ubiquitin ligase STUB1 and ubiquitin hydrolase USP7, were reported to promote or inhibit degradation of Foxp3 via ubiquitination, respectively (Chen et al., 2013; van Loosdregt et al., 2013). Finally, intracellular metabolism, and specifically the metabolic regulator mTOR (mammalian target of Rapamycin), has emerged as a key regulator of Foxp3 expression and Treg function. Early studies showed that weakened mTOR signaling leads to increased Foxp3 expression in iTregs in vitro (Delgoffe et al., 2009). However, recent studies using genetic models showed that complete ablation of mTOR in Tregs leads to compromised homeostasis and function of effector Tregs (Chapman et al., 2018; Sun et al., 2018). Despite these and other significant advances in understanding the molecular mechanisms regulating Foxp3, we lack a comprehensive picture of the regulatory networks that control Foxp3 expression.

In this study, we performed a genome-wide CRISPR/Cas9 knockout screen to identify the regulators of Foxp3 in mouse primary natural Treg cells. The screen results not only confirmed a number of known Foxp3 regulators but also revealed many novel factors that control Foxp3 expression. Gene ontology analysis showed that Foxp3 regulators are highly enriched in genes encoding subunits of the SAGA and SWI/SNF complexes, which we further validated by single gRNA knockout and flow cytometry analysis. The mammalian SWI/SNF complex is a multi-subunit complex with a core ATPase protein, either SMARCA4 (BRG1) or SMARCA2 (BRM), that uses energy derived from ATP hydrolysis to remodel nucleosomes on chromatin. Mouse genetic studies have demonstrated that conditional knockout of *Smarca4* leads to impaired differentiation of T lymphocytes (Gebuhr et al., 2003; Zhao et al., 1998). In addition, a previous report demonstrated that genetic deletion of *Smarca4* in Tregs using the Foxp3-Cre driver results in the development of a fatal inflammatory disorder reminiscent of Foxp3 mutant *scurfy* mice (Chaiyachati et al., 2013). The authors showed that while Treg development and Foxp3 expression was normal in *Smarca4* deficient Tregs, Treg function was nevertheless compromised due to impaired activation of TCR target genes, for example chemokine receptor genes in Tregs. This is consistent with the rapid association of SMARCA4-containing SWI/SNF complexes with chromatin following TCR activation in T cells (Zhao et al., 1998).

Biochemical studies have demonstrated that SMARCA4 is associated with both the canonical BAF complex (BAF) and Polybromo1-associated BAF complex (PBAF) (Xue et al., 2000; Yan et al., 2005). In addition, recent studies in embryonic stem cells (ESCs)(Gatchalian et al., 2018) and cancer cell lines (Alpsoy and Dykhuizen, 2018; Michel et al., 2018; Wang et al., 2019) have identified a BRD9-containing non-canonical complex or ncBAF complex (also referred to as GBAF or BRD9-containing BAF), which contains several shared subunits including SMARCA4, but is distinct from the BAF and PBAF complexes. Apart from uniquely incorporating BRD9, the ncBAF complex also contains GLTSCR1 or the paralog GLTSCR1L and lacks BAF- and PBAF-specific subunits ARID1A, ARID1B, ARID2, SMARCE1, SMARCB1, SMARCD2, SMARCD3, DPF1-3, PBRM1, BRD7, and PHF10. The distinct biochemical compositions of these three SWI/SNF complex assemblies suggest functional diversity. However, it is not known which SWI/SNF complex assemblies are expressed in Tregs and the potential roles of specific SWI/SNF variants in regulating Foxp3 expression and Treg development have not been studied in depth.

Here, we find that the BRD9-containing ncBAF complex promotes the expression of Foxp3, whereas the PBAF complex represses its expression. Furthermore, deletion of *Brd9* or PBAF component *Pbrm1* in Tregs results in reduced or enhanced suppressor activity, respectively, suggesting divergent regulatory roles of ncBAF and PBAF complexes in controlling Foxp3 expression and Treg function. Consistent with this model, we find that chemically-induced degradation of BRD9 by dBRD9 leads to reduced Foxp3 expression and compromised Treg function. Genome-wide binding studies revealed that BRD9 co-localizes with Foxp3, including at the CNS0 and CNS2 enhancers at the *Foxp3* locus. Furthermore, targeting BRD9 by sgRNA or dBRD9 reduces Foxp3 binding at the *Foxp3* locus and a subset of Foxp3 binding sites genome-wide, which results in differential expression of many Foxp3-dependent genes, indicating that BRD9 participates in the regulation of the Foxp3-dependent transcriptional program. Finally, we show that deletion of *Brd9* in Tregs reduced suppressor activity in an in vivo model of T cell transfer induced colitis, and improved anti-tumor immune responses in an MC38 colorectal cancer cell induced cancer model. In summary, we perform an unbiased genome-wide screen to identify genes and networks regulating Foxp3, and reveal ncBAF complex as a novel target that could be exploited to manipulate Treg function in vitro and in vivo.

## Results

### Genome-wide CRISPR screen in natural regulatory T cells identifies regulators of Foxp3

To screen for genes that regulate Foxp3 expression, we developed a pooled retroviral CRISPR sgRNA library by subcloning an optimized mouse genome-wide lentiviral CRISPR sgRNA library (lentiCRISPRv2-Brie) (Doench et al., 2016) into a newly engineered retroviral vector pSIRG-NGFR, which allowed us to efficiently transduce mouse primary T cells and to perform intracellular staining for Foxp3 without losing the transduction marker NGFR after cell permeabilization (Figure S1). Using this library, we performed CRISPR knockout screens on Tregs to identify genes that regulate Foxp3 expression. We activated CD4^+^Foxp3^+^ Tregs isolated from Rosa-Cas9/Foxp3^Thy1.1^ knock-in mice (Liston et al., 2008; Platt et al., 2014) with CD3 and CD28 antibodies and IL-2 (Figure 1A). Treg cells were transduced 24 hours post-activation with the pooled retroviral sgRNA library at multiplicity of infection of less than 0.2 to ensure only one sgRNA was transduced per cell. NGFR^+^ transduced Treg cells were collected on day 3 and day 6 to identify genes that are essential for cell proliferation and survival. In addition, the bottom quintile (NGFR^+^Foxp3^Low^) and top quintile (NGFR^+^Foxp3^High^) populations were collected on day 6 to identify genes that regulate Foxp3 expression. We validated the screen conditions by transducing Tregs with sgRNAs targeting *Foxp3* itself, as well as previously reported positive (*Cbfb*) (Rudra et al., 2009) and negative (*Dnmt1*) (Lal et al., 2009) regulators of Foxp3 (Figure 1B-D). Guide RNA sequences integrated within the genomic DNA of sorted cells were recovered by PCR amplification, constructed into amplicon libraries, and sequenced with a NextSeq sequencer.

**Figure 1.**
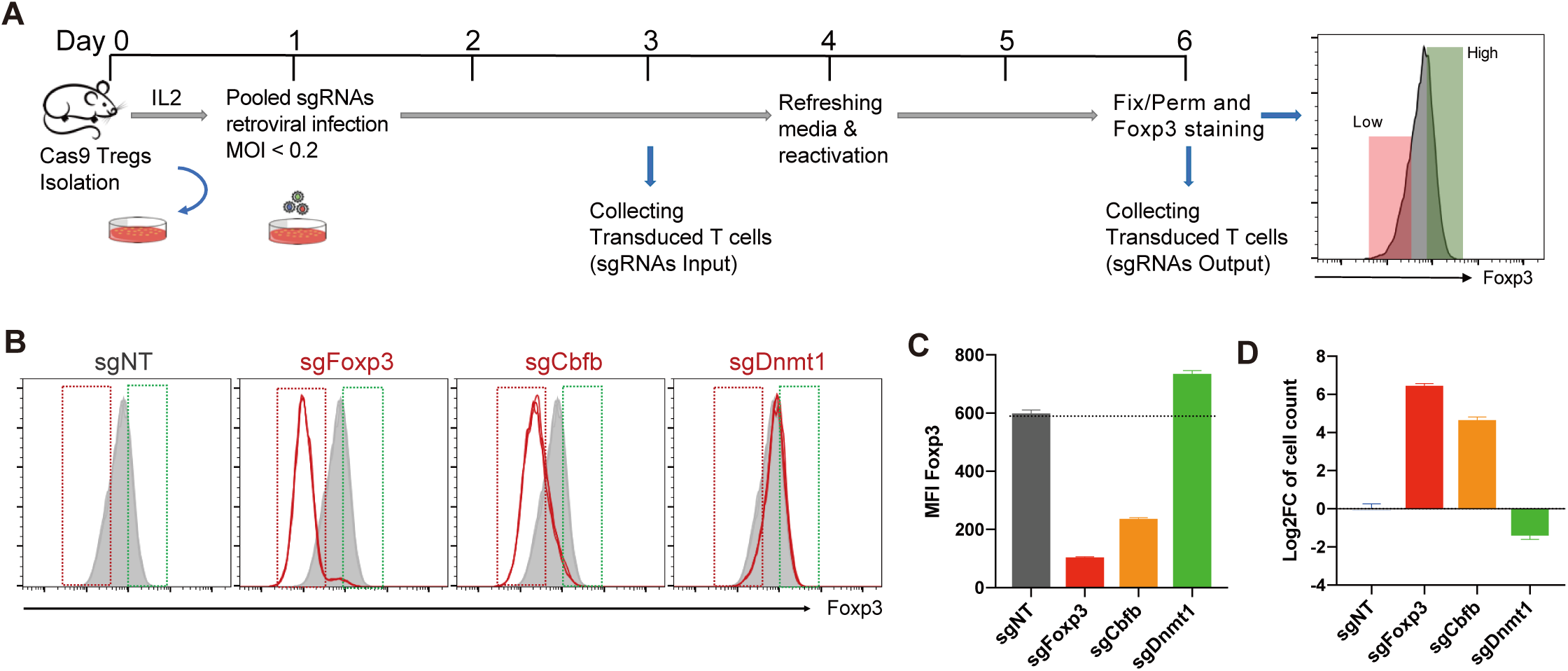
A genome-wide CRISPR screen in Treg cells. **A,** Workflow of the CRISPR screen in Tregs. **B-D,** Validation of the CRISPR screen conditions. **B,** FACS plots showing Foxp3 expression in Tregs after knocking out *Foxp3* (sgFoxp3), positive regulator *Cbfb* (sgCbfb), and negative regulator *Dnmt1* (sgDnmt1). Red and green gates were set based on Foxp3 low 20% and high 20% in sgNT Treg, respectively. **C,** Mean fluorescence intensity (MFI) of Foxp3 and **D,** Relative Log2FC of cell count comparing Foxp3^Low^ to Foxp3^High^ after deletion of the indicated target gene (n=3 per group). See also Figure S1 and S2.

The relative enrichment of sgRNAs between samples and hit identification were computed by MAGeCK, which generates a normalized sgRNA read count table for each sample, calculates the fold change of sgRNA read counts between two cell populations, and further aggregates information of four sgRNAs targeting each gene to generate a ranked gene list (Li et al., 2014a). Prior to hit calling, we evaluated the quality of screen samples by measuring the percentage of mapped reads to the sgRNA library and total read coverage, which showed a high mapping rate (79.8-83.4%) with an average of 236X coverage and a low number of missing sgRNAs (0.625-2.5%) (Figure S2). With the cutoff criteria of log2 fold change (LFC) >±0.5 and p-value less than 0.01, we identified 254 potential positive Foxp3 regulators enriched in the Foxp3^Low^ population and 490 potential negative Foxp3 regulators enriched in the Foxp3^High^ population (Figure 2A, 2B, and Table S1). In a parallel analysis, we also identified 22 and 1497 genes that affect cell expansion and contraction, respectively (p-value < 0.002, LFC>1, Figure S3 and Table S2). As expected, we identified genes belonging to pathways known to regulate Foxp3 expression both transcriptionally (*Cbfb*, *Runx3*) (Rudra et al., 2009) and post-transcriptionally through the regulation of Foxp3 protein stability (*Usp7, Stub1*) (Chen et al., 2013; van Loosdregt et al., 2013) (Figure 2C).

**Figure 2.**
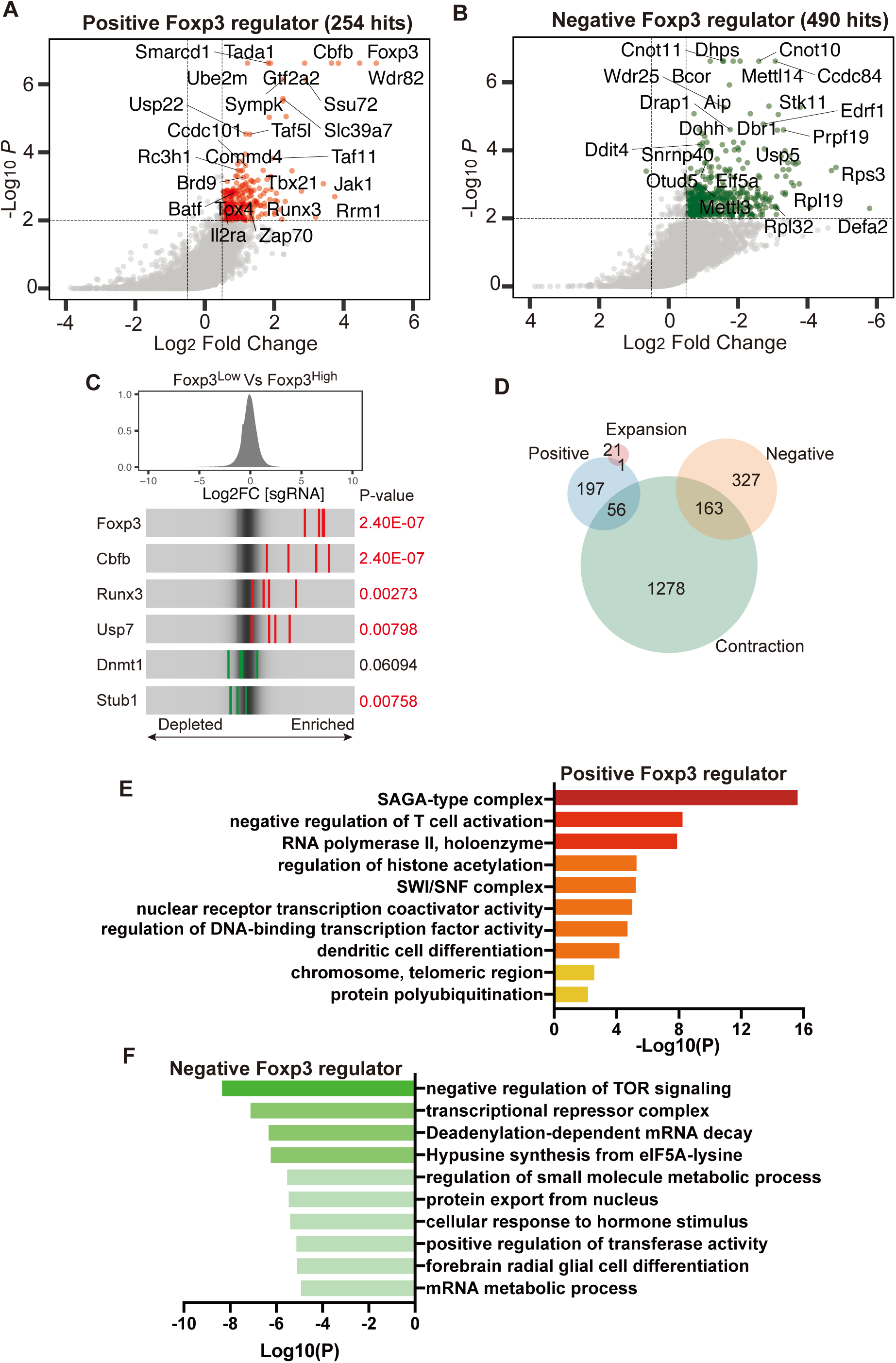
Identification of novel Foxp3 regulators in Treg cells. **A, B**, A scatter plot of the Treg screen result showing positive regulators (A) and negative regulators (B). Genes that have met cutoff criteria (P-value<0.01, and Log2FC >± 0.5) are shown as red dots for positive regulators and green dots for negative regulators. **C,** Distribution of sgRNA Log2FC comparing Foxp3^Low^ to Foxp3^High^. Red stripes represent sgRNAs from positive Foxp3 regulators, whereas green stripes represent sgRNAs from negative Foxp3 regulators. **D**, Venn diagram showing the overlap of Foxp3 regulators with genes involved in cell contraction or expansion. **E, F,** Gene ontology analysis of positive Foxp3 regulators **(E)** and negative Foxp3 regulators **(F)**. See also Figures S3 and S4; Tables S1, S2, S3, and S4.

We next compared the potential positive and negative regulators with genes involved in cell contraction and expansion to exclude hits that might affect Foxp3 expression indirectly by affecting cellular fitness in general, leaving 197 positive Foxp3 regulators and 327 negative Foxp3 regulators (Figure 2D and Table S3). Gene ontology analysis of positive Foxp3 regulators revealed a number of notable functional clusters including SAGA-type complex, negative regulation of T cell activation, RNA Polymerase II holoenzyme, positive regulation of histone modification, and SWI/SNF complex (Figure 2E, Table S4). Among negative Foxp3 regulators, genes are highly enriched in clusters related to negative regulation of TOR signaling, transcriptional repressor complex, mRNA decay and metabolism, and hypusine synthesis from eIF5A-lysine (Figure 2F, Table S4). Several of these pathways, including mTOR signaling, Foxp3 ubiquitination/deubiquitination, and transcriptional regulation, have been implicated in Foxp3 regulation previously, suggesting that our screen is robust for the validation of known pathways and the discovery of novel regulators of Foxp3. Among novel regulators, we identified many genes encoding subunits of the SAGA (*Ccdc101*, *Tada2b*, *Tada3, Usp22, Tada1, Taf6l, Supt5, Supt20*) and SWI/SNF (*Arid1a, Brd9*, *Smarcd1*) complexes (Table S4), strongly suggesting that these complexes could have indispensable roles for Foxp3 expression. We thus further validated and characterized the SAGA and SWI/SNF related complexes to understand their roles in Foxp3 expression and Treg function.

### Validation of the SAGA complex as a novel regulator of Foxp3 expression and Treg suppressor activity

The SAGA complex possesses histone acetyltransferase (HAT) and histone deubiquitinase (DUB) activity, and functions as a transcriptional co-activator through interactions with transcription factors and the general transcriptional machinery(Helmlinger and Tora, 2017; Koutelou et al., 2010). We identified *Ccdc101*, *Tada2b*, and *Tada3* in the HAT module, *Usp22* in the DUB module, and *Tada1*, *Taf6l*, *Supt5*, and *Supt20* from the core structural module among positive Foxp3 regulators that do not affect cell expansion or contraction (Figure S4A). We sought to validate the potential regulatory function of SAGA complex subunits by using sgRNAs to knock out individual subunits in Tregs and measure Foxp3 expression (Figure S4B, S4C). We found that deletion of every subunit tested resulted in a significant and 19-29% reduction in Foxp3 mean fluorescence intensity (MFI). We then further tested the function of SAGA subunit Usp22 in an in vitro suppression assay, which measures the suppression of T cell proliferation when conventional T cells are co-cultured with Tregs at increasing ratios. We found that Tregs transduced with sgRNAs targeting *Usp22* had compromised Treg suppressor activity compared with Tregs transduced with a non-targeting control sgRNA, with significantly more proliferation of T effector cells (Teff) at every ratio of Treg to Teff ratio tested (Figure S4D). These results provide independent validation of our genome-wide screen analyses for this class of chromatin regulators and demonstrate that the SAGA complex is essential for normal Foxp3 expression and that disrupting the SAGA complex by sgUsp22 reduces Treg suppressor function.

### Identification of the BRD9-containing ncBAF complex as a specific regulator of Foxp3 expression

We next wanted to characterize the role of SWI/SNF complex variants (BAF, ncBAF, and PBAF complexes) in Foxp3 expression. While these complexes share certain core subunits, such as the ATPase SMARCA4, each complex also contains specific subunits, for example the selective incorporation of the bromodomain containing protein BRD9 in ncBAF complexes (Figure 3A). Since the tissue-specific distribution and functional requirement for ncBAF complexes in primary T cells is not known, we performed co-immunoprecipitation assays to probe the composition of SWI/SNF-related complexes in Tregs. As expected, immunoprecipitation of SMARCA4, a core component of all three SWI/SNF complexes, revealed association of common subunits SMARCC1 and SMARCB1, as well as specific subunits ARID1A, BRD9, and PBRM1. Immunoprecipitations against ARID1A, BRD9, and PHF10 revealed the specific association of these subunits with BAF, ncBAF, and PBAF complexes, respectively (Figure 3A). These results established that all three SWI/SNF complexes are present with the expected composition in Tregs.

**Figure 3.**
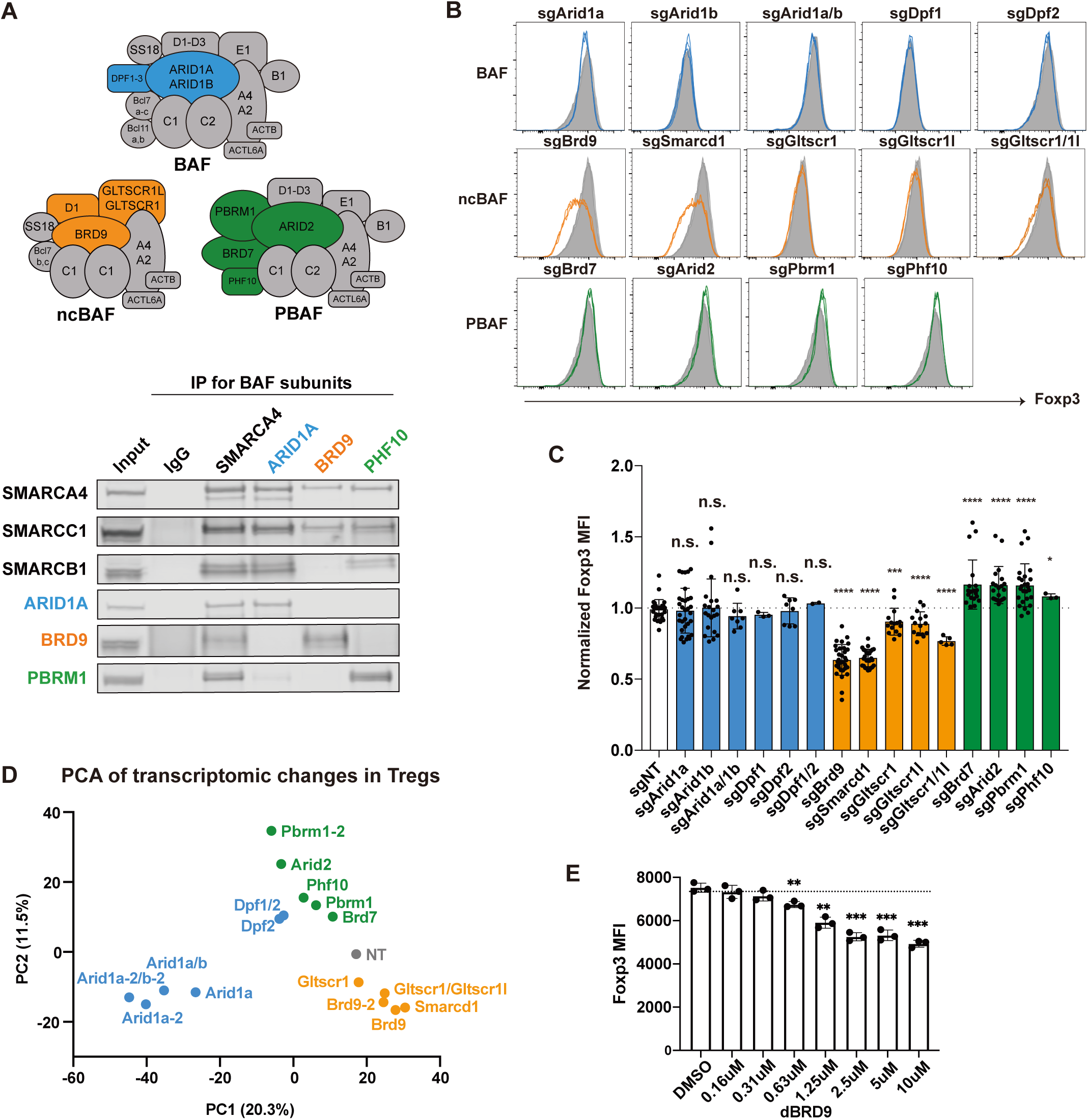
The three SWI/SNF complex assemblies have distinct regulatory roles for Foxp3 expression in Tregs. **A,** A diagram showing three different variants of SWI/SNF complexes: BAF, ncBAF, and PBAF. BAF-specific subunits (ARID1A, DPF1-3) are colored blue, ncBAF-specific subunits (BRD9, SMARCD1, GLTSCR1L, GLTSCR1) colored orange, and PBAF-specific subunit (PBRM1, ARID2, BRD7, PHF10) colored green. Shared components among complexes are colored gray. Immunoprecipitation assay of ARID1A, BRD9, and PHF10, and BRG1 in Tregs. The co-precipitated proteins were probed for shared subunits (SMARCA4, SMARCC1, SMARCB1), BAF-specific ARID1A, ncBAF-specific BRD9, and PBAF-specific PBRM1. **B,** FACS histogram of Foxp3 expression in Tregs after sgRNA knockout of the indicated SWI/SNF subunits. **C,** Mean fluorescence intensity (MFI) of Foxp3 after sgRNA knockout of the indicated SWI/SNF subunits. Data represents mean and standard deviation of biological replicates (n = 3-21). **D,** Principal component analysis of RNA-seq data collected from Tregs transduced with guides against the indicated SWI/SNF subunits. In cases where two independent guides were used to knockdown a gene, the second guide for targeting gene indicated as “-2”. **E,** MFI of Foxp3 expression in Tregs after treatment with either DMSO or 0.16-10 μM dBRD9 for 4 days. Data represent mean ± s.d. Statistical analyses were performed using unpaired two-tailed Student’s t test (ns: p≥0.05, *p<0.05, **p<0.01, ***p<0.001, ****p<0.0001).

In our screen, we identified *Brd9*, *Smarcd1, Arid1a* and *Actl6a* among positive regulators of Foxp3, whereas SWI/SNF shared subunits *Smarca4*, *Smarcb1*, *Smarce1*, and *Actl6a* were identified in cell contraction (Table S3). This suggests a potential regulatory role for ncBAF and/or BAF complexes. To explore the specific function of BAF, ncBAF, and PBAF complexes in Foxp3 expression, we cloned independent sgRNAs to knockout unique subunits for each complex, and measured Foxp3 MFI in sgRNA transduced Tregs. We observed an essential role for the ncBAF complex in Foxp3 expression in Tregs. Specifically, knockdown of ncBAF specific subunits, including *Brd9* and *Smarcd1*, significantly diminished Foxp3 expression by nearly 40% in Tregs (Figure 3B, 3C). Knockdown of ncBAF-specific paralogs *Gltscr1* and *Gltscr1l* individually resulted in a slight reduction in Foxp3 expression, which was further reduced in the *Gltscr1*/*Gltscr1l* double knockout, suggesting that these two paralogs can compensate in the regulation of Foxp3 expression (Figure 3C). In contrast, knockdown of PBAF specific subunits, including *Pbrm1*, *Arid2*, *Brd7,* and *Phf10*, significantly enhanced Foxp3 expression by as much as 17% (Figure 3C, green). Knockdown of BAF specific subunits *Arid1a, Arid1b, Dpf1*, or *Dpf2* did not significantly affect Foxp3 expression (Figure 3C, blue). To determine if ARID1A and ARID1B could be compensating for one another, we performed *Arid1a*/*Arid1b* double deletion and found that deletion of either or both ARID paralogs resulted in slight, but non-significant reduction in Foxp3 MFI (Figure 3C, blue). These data suggest that ncBAF and PBAF have opposing roles in the regulation of Foxp3 expression. To further explore the role of different SWI/SNF complexes in Treg genome-wide transcription, we performed RNA sequencing from Tregs with knockdown of variant-specific subunits with one or two independent guide RNAs and conducted principal component analysis, which showed that the ncBAF, PBAF, and BAF also have distinct effects at whole transcriptome level in Tregs (Figure 3D).

We then made use of a recently developed chemical BRD9 protein degrader (dBRD9)(Remillard et al., 2017) as an orthogonal method to probe BRD9 function. dBRD9 is a bifunctional molecule that links a small molecule that specifically binds to the bromodomain of BRD9 and another ligand that recruits the cereblon E3 ubiquitin ligase. We confirmed that treatment of Tregs with dBRD9 resulted in reduced BRD9 protein levels (Figure S5A). Similar to sgRNA depletion of *Brd9*, dBRD9 treatment significantly decreased Foxp3 expression in Treg cells in a concentration-dependent manner, without affecting cell viability or proliferation (Figure 3E, Figure S5B). These data demonstrate the requirement for BRD9 in maintenance of Foxp3 expression using both genetic and chemically-induced proteolysis methods.

### BRD9 regulates Foxp3 binding at the CNS0 and CNS2 enhancers and a subset of Foxp3 target sites

To dissect the molecular mechanism of how ncBAF and PBAF complexes regulate Foxp3 expression in Treg cells, we performed chromatin immunoprecipitation followed by genome-wide sequencing (ChIP-seq) in Tregs using antibodies against the ncBAF-specific subunit BRD9, the PBAF-specific subunit PHF10 and the shared enzymatic subunit SMARCA4. Data generated from these ChIP-seq experiments revealed that BRD9, SMARCA4, and PHF10 co-localize at CNS2 in the *Foxp3* gene locus and at CNS0 found within the *Ppp1r3f* gene immediately upstream of *Foxp3* (Figure 4A). Since CNS2 was previously shown to regulate stable Foxp3 expression through a positive feedback loop involving Foxp3 binding(Feng et al., 2014; Li et al., 2014b), and Foxp3 is additionally bound at CNS0 in Tregs(Kitagawa et al., 2017), we rationalized that ncBAF and/or PBAF complexes might affect Foxp3 expression by regulating Foxp3 binding at CNS2/CNS0. We therefore performed Foxp3 ChIP-seq in Tregs transduced with sgNT, sgFoxp3, sgBrd9 or sgPbrm1. We observed a dramatic reduction in Foxp3 binding at CNS2/CNS0 in sgFoxp3 transduced cells, as expected, and there was also marked reduction of Foxp3 binding at CNS2/CNS0 in *Brd9*-depleted Tregs (Figure 4A). In contrast, we observed a subtle increase in Foxp3 binding at CNS2/CNS0 in *Pbrm1* sgRNA transduced Tregs, which could explain why *Pbrm1* emerged as a negative regulator of Foxp3 expression in our validation studies (Figure 4A). These data suggest that BRD9 positively regulates Foxp3 expression by promoting Foxp3 binding to its own enhancers.

**Figure 4.**
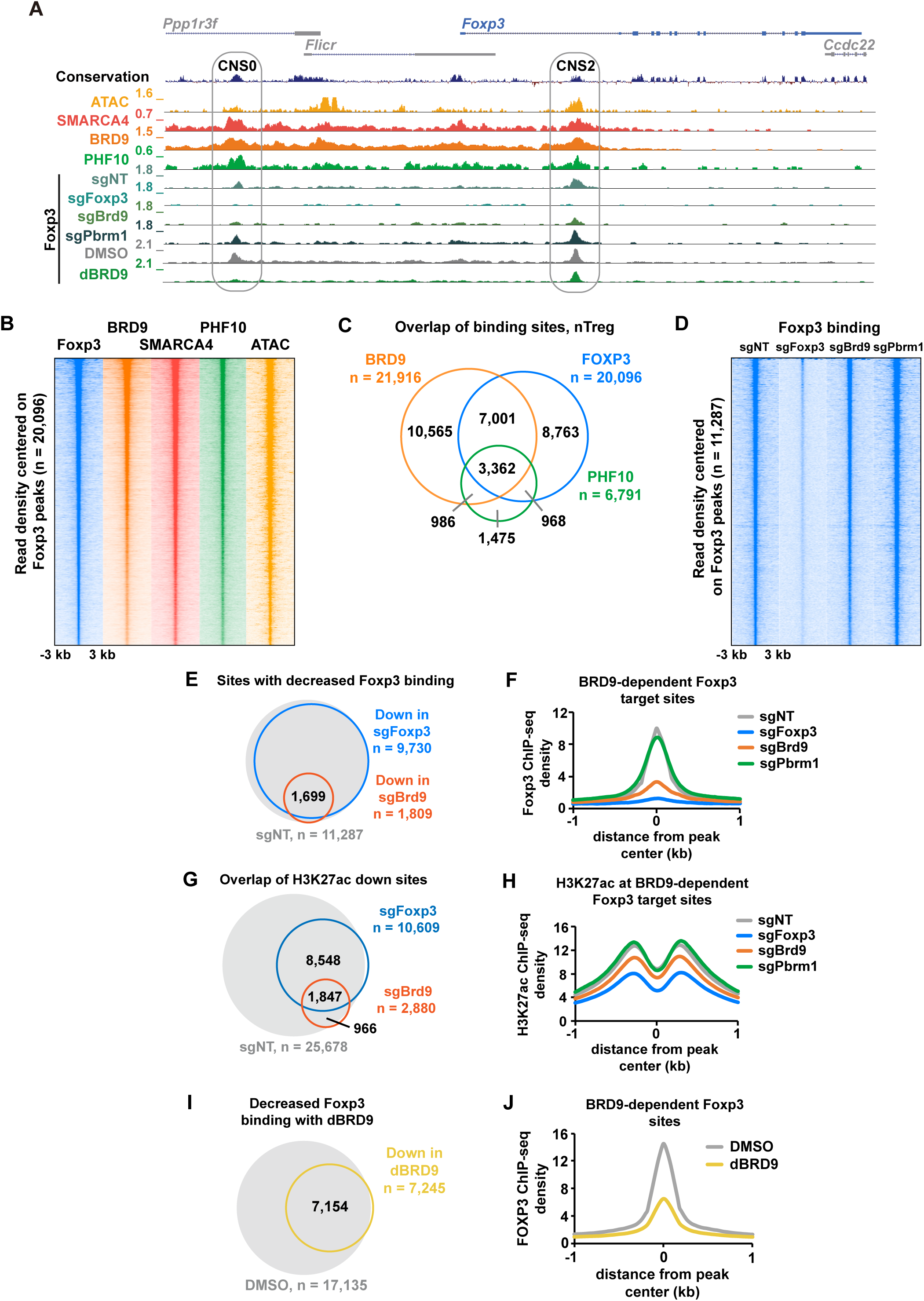
BRD9 deletion reduces Foxp3 binding at CNS0, CNS2 enhancers and a subset of Foxp3 target sites. **A,** Genome browser tracks of SMARCA4, BRD9, PHF10 ChIP-seq and ATAC-seq signal, as well as Foxp3 ChIP-seq in sgNT, sgFoxp3, sgBrd9 and sgPbrm1 Tregs and Foxp3 in DMSO and dBRD9 treated Tregs (2.5 μM dBRD9 for 4 days). *Foxp3* locus is shown with CNS0 and CNS2 enhancers indicated in gray ovals. **B**, Heat map of Foxp3, BRD9, SMARCA4, PHF10 ChIP-seq and ATAC-seq signal ± 3 kb centered on Foxp3-bound sites in Tregs, ranked according to Foxp3 read density. **C,** Venn diagram of the overlap between ChIP-seq peaks in Tregs for BRD9, Foxp3, and PHF10 (hypergeometric p value of BRD9:Foxp3 overlap = e^-30704^, hypergeometric p value of PHF10:Foxp3 overlap = e^-13182^, hypergeometric p value of BRD9:PHF10 overlap = e^-12895^). **D,** Heat map of Foxp3 ChIP-seq signal in sgNT, sgFoxp3, sgBrd9 and sgPbrm1 Tregs ± 3 kilobases (kb) centered on Foxp3-bound sites in sgNT, ranked according to read density. **E**, Venn diagram of the overlap (hypergeometric p value = e^-11,653^) between sites that significantly lose Foxp3 binding (FC 1.5, Poisson p value < 0.0001) in sgFoxp3 and sgBrd9, overlaid on all Foxp3-bound sites in sgNT (in gray). **F**, Histogram of Foxp3 ChIP read density ± 1 kb surrounding the peak center of sites that significantly lose Foxp3 binding in both sgFoxp3 and sgBrd9 (n=1,699) in sgNT, sgFoxp3, sgBrd9 and sgPbrm1. **G,** As in **E**, but for sites that lose H3K27ac (FC 1.5, Poisson p value < 0.0001, hypergeometric p value of overlap = e^-7,938^). **H,** As in **F**, but for H3K27ac ChIP read density. **I,** As in **E**, but for sites that significantly lose Foxp3 binding in dBRD9 treated Tregs versus DMSO (FC 1.5, Poisson p value < 0.0001). **J**, As in **F**, but for DMSO and dBRD9 treated cells.

We then extended this analysis to examine the cooperation between BRD9 and Foxp3 genome-wide. Notably, we find co-binding of BRD9, SMARCA4, and PHF10 with Foxp3 at a subset of Foxp3-bound sites (Figure 4B, 4C). All four factors localize to promoters, intronic, and intergenic regions of the genome and their binding correlates well with chromatin accessibility as measured by assay of transposase-accessible chromatin with sequencing (ATAC-seq) (Figure 4B, S6A). Motif analysis of Foxp3-bound sites revealed an enrichment for motifs recognized by ETS and RUNX transcription factors consistent with what has been previously shown(Samstein et al., 2012). ETS and RUNX motifs were also among the most significant motifs at both BRD9-bound sites, along with an enrichment of the CTCF motif as we and others previously reported(Gatchalian et al., 2018; Michel et al., 2018) (Figure S6B). These results demonstrate that ncBAF and PBAF complexes are co-localized with Foxp3 at Foxp3 binding sites genome-wide.

To assess the requirement for BRD9 or PBRM1 in Foxp3 targeting genome-wide, we analyzed Foxp3 binding in Tregs transduced with sgNT, sgFoxp3, sgBrd9, or sgPbrm1 at all Foxp3 binding sites (Figure 4D). As expected, we find that Foxp3 binding is lost at over 85% of its binding sites in sgFoxp3-transduced Treg cells (Figure 4E). Foxp3 binding at a subset of these sites is also significantly reduced in sgBrd9-transduced Tregs (FC 1.5, Poisson p < 0.0001), suggesting that BRD9 is required for Foxp3 binding at a subset of its target sites (Figure 4E). This is a specific function of BRD9, as Foxp3 binding does not change in *Pbrm1*-depleted Tregs at these BRD9-dependent sites (Figure 4F). ChIP-seq for the active histone mark H3 lysine27 acetylation (H3K27ac) revealed that BRD9 and Foxp3 cooperate to maintain H3K27ac at over 1,800 shared sites (Figure 4G). At BRD9-dependent Foxp3 sites, for example, we observed a reduction in H3K27ac in sgFoxp3 and sgBrd9-transduced Tregs, but not in sgPbrm1-transduced Tregs (Figure 4H). Using dBRD9, we confirmed that BRD9 binding to chromatin is reduced (Figure S6C). We further recapitulated our observation that BRD9 loss results in diminished Foxp3 binding to chromatin at a subset of Foxp3 target sites (Figure 4I, 4J, S6D), including at CNS2 and CNS0 (Figure 4A). These data demonstrate that BRD9 co-binds with Foxp3 at the *Foxp3* locus to positively reinforce its expression. BRD9 additionally promotes Foxp3 binding and H3K27ac levels at a subset of Foxp3 target sites both by potentiating Foxp3 expression and through direct epigenetic regulation at BRD9/Foxp3 co-bound sites.

### BRD9 co-regulates the expression of Foxp3 and a subset of Foxp3 target genes

Based on co-binding of BRD9 and Foxp3 at Foxp3 target sites, we assessed the effects of BRD9 ablation on the transcription of Foxp3 target genes. We performed RNA-seq in Tregs transduced with sgFoxp3, sgBrd9, or sgNT. Consistent with Foxp3’s role as both transcriptional activator and repressor, we observed down-regulation and up-regulation of 793 and 532 genes, respectively, in *Foxp3* sgRNA transduced Tregs, which are enriched in ‘cytokine production’, ‘regulation of defense response’, and ‘regulation of cell adhesion’ (Figure 5A, 5B). Of these, 67% are directly bound by Foxp3 in our ChIP-seq dataset and 60% are co-bound by Foxp3 and BRD9 (Figure 5C). Deletion of BRD9 resulted in transcriptional changes that strongly correlated with the transcriptional changes in sgFoxp3 transduced Tregs (r^2^ = 0.534, Linear regression analysis; Figure 5D). Indeed, gene set enrichment analysis (GSEA) demonstrated that the sgBrd9 up-regulated genes are significantly enriched among genes that increase upon Foxp3 knockdown, while the sgBrd9 down-regulated genes are enriched among genes that decrease in sgFoxp3 Tregs (Figure 5E). We also performed RNA-seq for Tregs treated with either vehicle or the dBRD9 degrader and observed a similar significant enrichment for dBRD9 affected genes among the Foxp3 up- and down-regulated genes (Figure 5F). Notably, the BRD9-dependent target gene sets generated from our RNA-seq data were among the most significantly enriched dataset of 9,229 immunological, gene ontology and curated gene sets when analyzed against the sgFoxp3 transduced Treg expression data (Figure 5G). In addition, both datasets were significantly enriched for genes that are differentially expressed between Tregs and conventional T cells(Feuerer et al., 2010), and between Foxp3 mutant Tregs from *scurfy* mice and wild-type Tregs(Hill et al., 2007). These data define a role for BRD9 in Tregs through specifically regulating the expression of Foxp3 itself and a subset of Foxp3 target genes.

**Figure 5.**
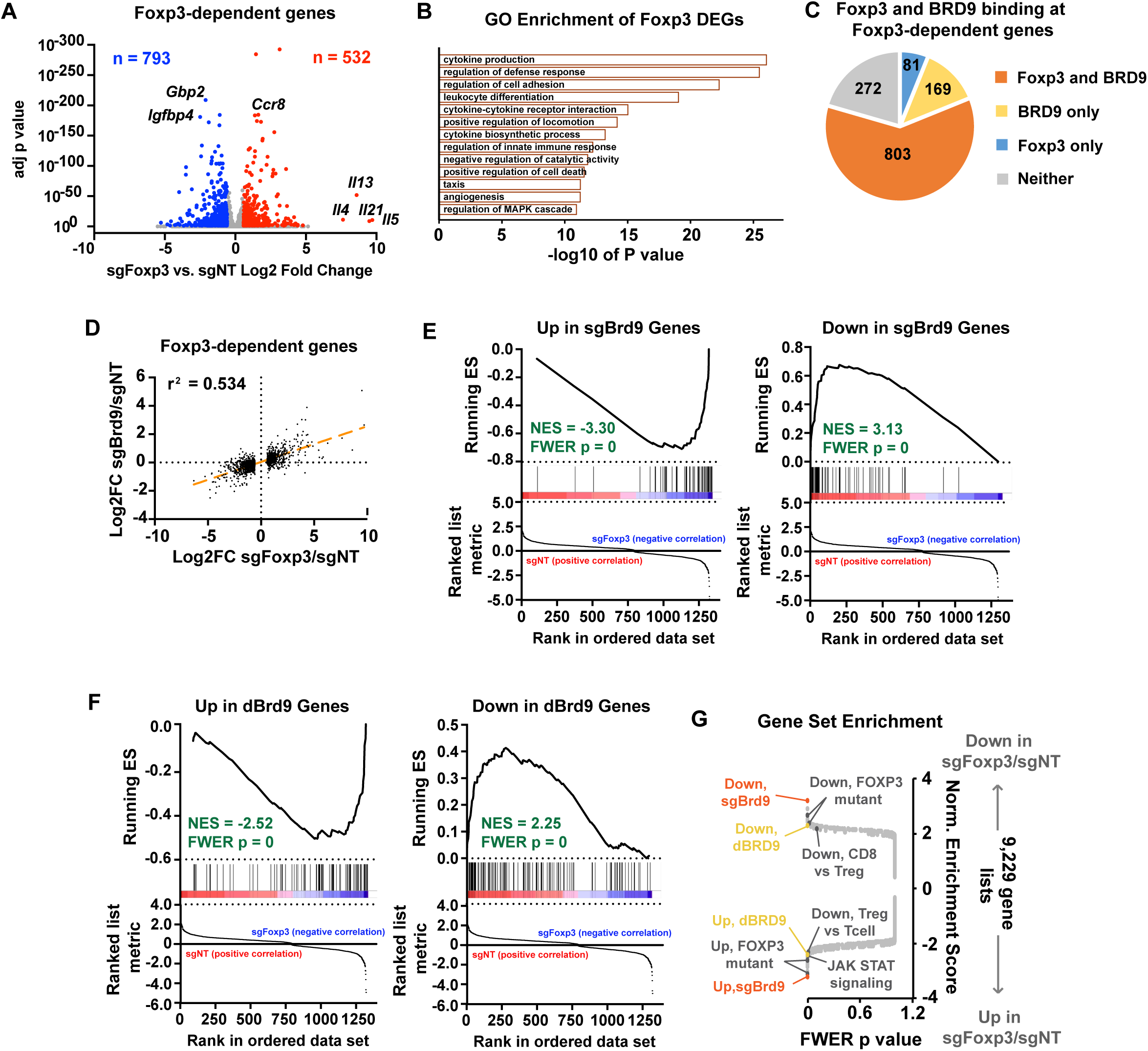
BRD9 co-regulates the expression of Foxp3 and a subset of Foxp3 target genes. **A,** Volcano plot of log2 fold change RNA expression in sgFoxp3/sgNT Tregs versus adjusted p value (Benjamin-Hochberg). Number of down- and up-regulated genes are indicated, which are colored blue and red, respectively. **B,** Significance of enrichment of Foxp3-dependent genes in each gene ontology. **C,** Pie chart of Foxp3 and BRD9 binding by ChIP-seq for Foxp3-dependent genes. **D,** Scatterplot of the mRNA log2 fold changes in sgFoxp3/sgNT and sgBrd9/sgNT for Foxp3-dependent genes. Linear regression analysis was performed to calculate the r^2^. Best fit is represented as an orange dashed line. **E,** Gene set enrichment analysis (GSEA) enrichment plot for up- and down-regulated genes in sgBrd9/sgNT compared with RNA-seq data of genes that significantly change in sgFoxp3/sgNT Tregs. ES: Enrichment Score, NES: Normalized Enrichment Score, FWER: Familywise Error Rate. **F**, As in **E**, but for up- and down-regulated genes in dBRD9/DMSO Tregs. **G,** GSEA of the sgFoxp3/sgNT RNA-seq data; plot shows the familywise error rate (FWER) p value versus the normalized enrichment score (NES). See also Table S5.

### ncBAF complex is required for normal Treg suppressor activity in vitro and in vivo

The divergent roles of ncBAF and PBAF complexes in regulating Foxp3 expression suggested that these complexes might also differentially affect Treg suppressor function. We performed sgRNA knockdown of ncBAF-specific *Brd9* and *Smarcd1* or PBAF-specific *Pbrm1* and *Phf10* in Tregs and measured their function by conducting an *in vitro* suppression assay. Tregs depleted of *Brd9* or *Smarcd1* exhibited significantly reduced suppressor function, whereas depletion of *Pbrm1* or *Phf10* resulted in significantly enhanced suppressor function (Figure 6A, S7A). These data demonstrate that the opposing regulation of Foxp3 expression by ncBAF and PBAF complexes results in decreased/increased Treg suppressor activity upon ncBAF or PBAF subunit deletion, respectively. Similar to sgRNA depletion of *Brd9*, Tregs treated with dBRD9 also showed significantly and specifically compromised Treg suppressor function *in vitro* (Figure S7B). These results underscore the requirement for BRD9 in Foxp3 expression maintenance and Treg suppressor activity, and further demonstrate that dBRD9 reduces Treg suppressor activity without impairing T effector responses in vitro.

**Figure 6.**
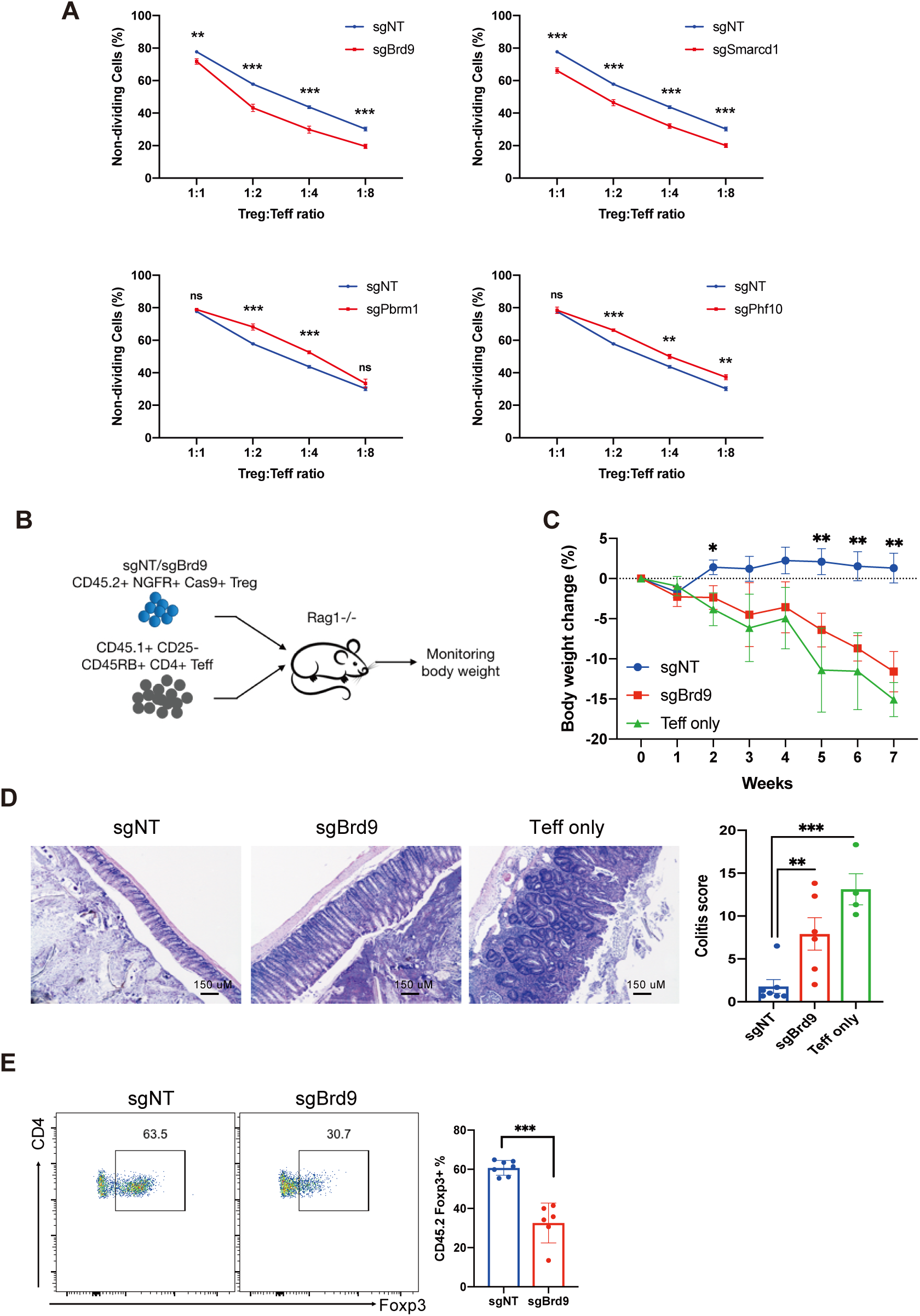
The ncBAF complex regulates Treg suppressor function in vitro and in vivo. **A.** In vitro suppression assay of Tregs with sgRNA knockout of *Brd9, Smarcd1, Pbrm1,* and *Phf10* (n=3 per group, data represent ± s.d.). sgNT was used as non-targeting control. **B-F.** Experiment to measure Treg function of sgNT or sgBrd9 knockout Treg cells relative to no Tregs in a T cell transfer induced colitis model. **B,** Experimental procedure. **C,** Body weight loss. **D,** Colon histology (left) and colitis scores (right). **E,** Percentage of Foxp3+ cells in transferred CD45.2+CD4+ Treg population at end point. (n=4-6 per group. Data represent mean ± s.e.m.) Statistical analyses were performed using unpaired two-tailed Student’s t test (ns: p≥0.05, *p<0.05, **p<0.01, ***p<0.001).

To test if BRD9 also affects Treg function *in vivo*, we utilized a T cell transfer-induced colitis model. In this model, *Rag1* knockout mice were either transferred with CD45.1^+^ CD4^+^ CD25^-^CD45RB^High^ effector T cell (Teff) only, or co-transferred with Teff along with CD45.2^+^ Tregs transduced with *Brd9* sgRNA (sgBrd9) or control sgRNA (sgNT) (Figure 6B). Mice transferred with Teff cells alone lost body weight progressively due to development of colitis. Co-transfer of Tregs transduced with sgNT protected recipient mice from weight loss, whereas co-transfer of sgBrd9 transduced Tregs failed to protect recipients from losing weight (Figure 6C). The mice transferred with *Brd9*-depleted Tregs showed significant colitis pathology at seven weeks compared to mice that received control Tregs (Figure 6D). Furthermore, *Brd9* depletion also led to compromised Treg stability after transfer, manifested by reduced Foxp3^+^ cell frequencies within the CD45.2^+^CD4^+^ transferred Treg population (Figure 6E). These results demonstrate that BRD9 is an essential regulator of normal Foxp3 expression and Treg function in a model of inflammatory bowel disease *in vivo*.

In addition to their beneficial role in preventing autoimmune diseases, Tregs have also been shown to be a barrier to anti-tumor immunity. We therefore wondered whether we could exploit the compromised suppressor function shown in *Brd9* deficient Tregs to disrupt Treg-mediated immune suppression in tumors. We used the MC38 colorectal tumor cell line to induce cancer due to the prominent role Tregs play in this cancer model(Delgoffe et al., 2013). *Rag1* knockout mice were used as recipients for adoptive transfer of Treg depleted-CD4 and CD8 T cells (Teff) only, or co-transfer of Teff with Tregs transduced with either sgBrd9 or control sgNT. MC38 tumor cells were implanted subcutaneously on the following day (Figure 7A). Transfer of sgNT Tregs allowed for significantly faster tumor growth compared to mice that received Teff cells only (“No Treg”) due to suppression of the anti-tumor immune response by Tregs (Figure 7B, 7C).

**Figure 7.**
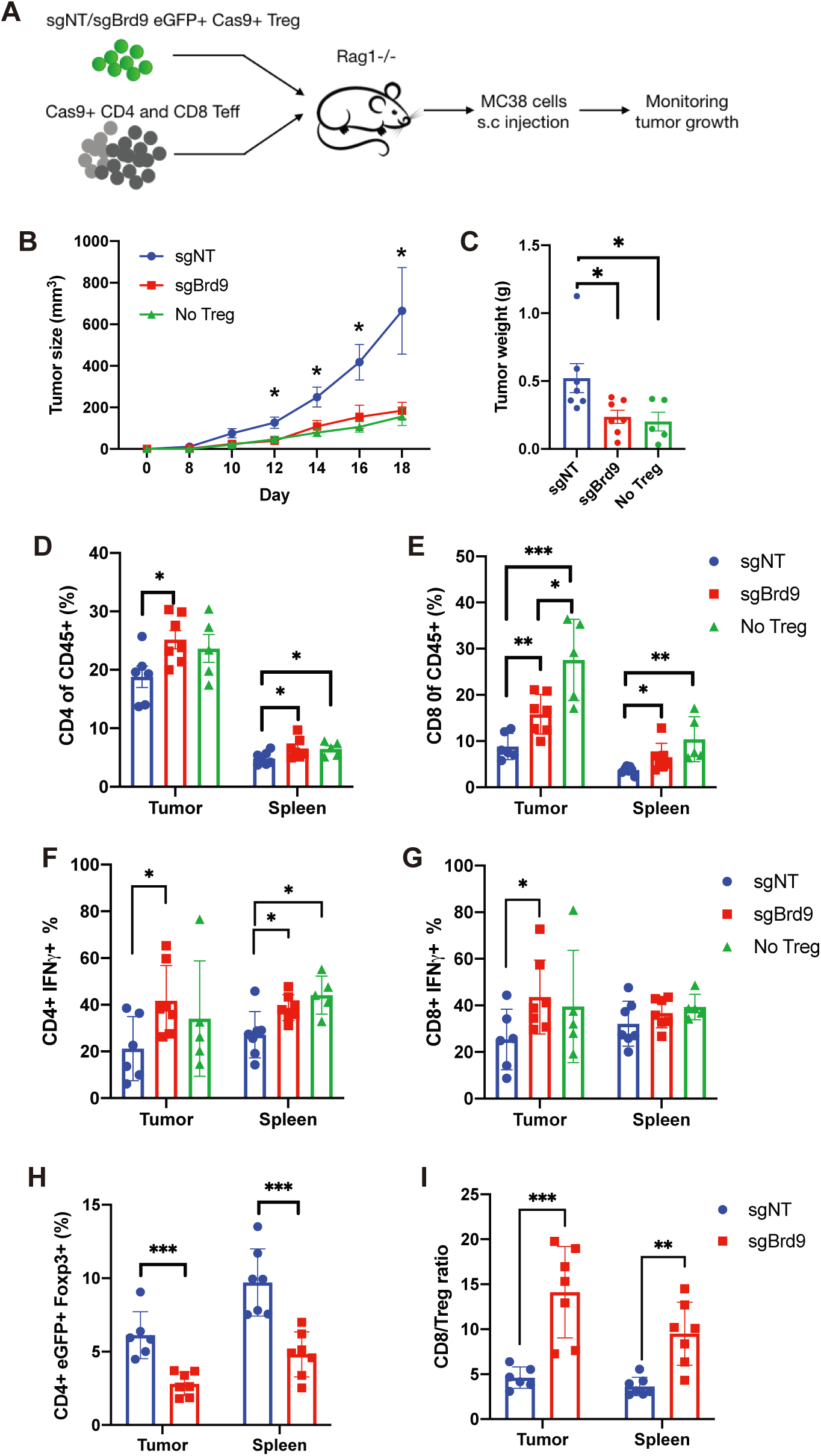
Targeting BRD9 in Treg improves anti-tumor immunity. **A,** Experiment procedure to measure function of sgNT or sgBrd9 knockout Treg cells relative to no Tregs in MC38 tumor model. **B**, Tumor growth curve. **C**, Tumor weight at end point. **D,E**, Bar graph of total CD4 T cells (**D**) and CD8 T cells (**E**) percentage in CD45+ immune cell population. **F,G**, Bar graph of IFN-γ+ cell percentage in CD4 T cells (**F**) and in CD8 T cells (**G**). **H**, Bar graph of CD4+eGFP+Foxp3+ donor cells in CD4+ T cells**. I,** Ratio of CD8/Treg. (n=5-7 per group. Data represent mean ± s.e.m.) Statistical analyses were performed using unpaired two-tailed Student’s t test (ns: p≥0.05, *p<0.05, **p<0.01, ***p<0.001).

Furthermore, tumor growth in mice that received sgBrd9 transduced Tregs was significantly slower than mice that received sgNT Tregs consistent with our findings that Brd9 knockdown reduces Treg suppressor activity (Figure 7B, 7C). Both CD4 and CD8 T cell tumor infiltration significantly increased in mice that received sgBrd9 transduced Tregs compared to sgNT Tregs (Figure 7D, 7E). Additionally, the percent of IFN-γ producing intra-tumor CD4 and CD8 T cells in mice that received sgBrd9 transduced Tregs was significantly greater than the sgNT Treg condition, and comparable to the transfer of Teff alone (“No Treg”) (Figure 7F, 7G). Consistent with our findings that BRD9 is required for Treg persistence in vivo (Figure 6E), the percentage of transferred Treg cells was reduced in mice that received sgBrd9 transduced Tregs relative to sgNT Tregs (Figure 7H). Overall, a 2-3 fold increase in the ratio of CD8 T cells to Tregs in tumor and spleen was observed in the sgBrd9 versus the sgNT condition, consistent with the enhanced anti-tumor immune response in mice that received sgBrd9 transduced Tregs (Figure 7I). This experiment demonstrates that BRD9 promotes stable Treg function in MC38 tumors and knockdown of Brd9 in Tregs improves anti-tumor immunity in this context.

## Discussion

In this study, we performed a genome-wide CRISPR screen to identify positive and negative regulators of Foxp3 expression in mouse natural Tregs. Among positive regulators, we identified *Cbfb and Runx3*, consistent with previous reports showing a requirement for CBF-β/Runx3 in Foxp3 expression and Foxp3-dependent target gene expression(Kitoh et al., 2009; Rudra et al., 2009). Among the novel positive regulators, we discovered subunits from two chromatin remodeling complexes, the BRD9-containing ncBAF and SAGA complexes. Independent validation and functional assays demonstrated an essential role for the ncBAF complex and SAGA complex in Foxp3 expression and Treg suppressor function.

Our screens also confirmed several known negative regulators of Foxp3, including DNA methyl-transferase *Dnmt1* and the ubiquitin ligase *Stub1*. Additionally, we identified multiple negative regulators of the mTOR pathway as Foxp3 negative regulators (*Tsc2*, *Flcn*, *Ddit4*, *Sesn2*, *Nprl2*), confirming an essential role for mTOR in homeostasis and function of activated Tregs(Chapman et al., 2018; Sun et al., 2018). Among novel negative Foxp3 regulators, we uncovered genes encoding regulators of RNA metabolism, which have no previously reported function in Foxp3 expression. For example, *Mettl3* and *Mettl14* form a methyltransferase complex that is essential for the m^6^A methylation of RNA, which is recognized as an important regulatory mechanism for a wide range of biological processes, including RNA stability, protein translation, stem cell self-renewal, cell lineage determination, and oncogenesis(Yue et al., 2015). Our screen suggests a potentially novel role for RNA m^6^A methylation in post-transcriptional regulation of Foxp3. Together, our genome-wide screens provide the first comprehensive picture of the complex regulatory network controlling Foxp3 expression levels, and reveal previously unknown pathways and factors that warrant further investigation.

Following the identification of SWI/SNF subunit genes among Foxp3 regulators, we endeavored to characterize the roles of the three SWI/SNF-related complexes by deleting subunits unique to each of the ncBAF, BAF, and PBAF complexes. We observed specific and divergent roles of ncBAF and PBAF complexes in regulating Foxp3 expression in Tregs. In contrast, deletion of BAF-specific subunits had a slight, but non-significant effect on Foxp3 expression. Nevertheless, several SWI/SNF core subunits were recovered in our screen among genes that regulate Treg cell contraction, suggesting that BAF complexes may regulate Treg activation or proliferation in response to TCR stimulation used to activate and culture Tregs in our screen. This is consistent with a role for *Smarca4* in Treg activation and control of autoimmunity *in vivo* independent of affecting Foxp3 expression, which is not changed in Foxp3-Cre:Smarca4^f/f^ Tregs(Chaiyachati et al., 2013). Thus, deletion of *Smarca4* or other BAF complex subunits likely results in overall defects in Treg fitness, whereas deletion of ncBAF subunits appears to have a selective effect on Foxp3 expression and its target genes. Mechanistically, we find that the ncBAF complex co-binds and cooperates with Foxp3 to potentiate its binding to the CNS2 and CNS0, enhancers of the *Foxp3* locus. In addition to the Foxp3 locus itself, our ChIP-seq analysis revealed that ncBAF also binds to regulatory elements in a subset of Foxp3 target genes to regulate their gene expression. One possibility is that reduced Foxp3 expression results in lowered Foxp3 binding at a select group of target genes, but what differentiates the dosage-dependent binding of Foxp3 at these sites compared to other unaffected sites remains unclear.

Finally, we tested the vivo relevance of our findings by disrupting the ncBAF subunit Brd9 in Tregs in mouse models of inflammatory bowel disease and cancer. Knockdown of *Brd9* in Tregs weakened their suppressor function in a model of T cell induced colitis, leading to exacerbated disease progression. In the context of cancer, we found that transfer of *Brd9* deficient Tregs failed to restrict anti-tumor immune responses in the MC38 cell induced cancer model, leading to slower tumor growth. Currently, there is a concerted effort to develop compounds targeting a number of SWI/SNF complex subunits to modulate their function. Our data show that bromodomain-directed degradation of BRD9 by dBRD9 recapitulates the effects of *Brd9* genetic deletion, suggesting that the ncBAF complex can be targeted with small molecules to control Foxp3 expression and Treg function. Thus, through the unbiased screen of Foxp3 regulators, we have identified novel proteins that can potentially be targeted to manipulate Treg homeostasis and function in autoimmune diseases and cancer.

## Acknowledgments

We would like to thank C. Gordon for mouse colony management, B. Kuo and X. Hu for assistance in plasmid extraction and mouse dissection, N. Hah and G. Chou for assistance in RNA-seq, ChIP-seq, CRISPR screen experiments, E. Shifrut and A. Marson (UCSF) for providing us the R script to generate the sgRNA distribution histograms, A. Williams and M. Shokhirov for bioinformatic assistance, and M. Downes and R. M. Evans for helpful discussion. C.S.L. was partly supported by the Albert G. and Olive H. Schlink Foundation. J.G. was supported by the Salk Institute T32 Cancer Training Grant T32CA009370 and the NIGMS NRSA F32 GM128377-01. D.C.H. was supported by the National Institutes of Health (GM128943-01, CA184043-03), the V Foundation for Cancer Research V2016-006, and the Leona M. and Harry B. Helmsley Charitable Trust. Y.Z. was supported by the NOMIS Foundation, the Crohn’s and Colitis Foundation, the Leona M. and Harry B. Helmsley Charitable Trust, and National Institute of Health (AI107027 and OD023689). This work was also supported by National Cancer Institute funded Salk Institute Cancer core facilities (CA014195).

## Author Contributions

Conceptualization: C.S.L., J.G., D.C.H. and Y.Z. Methodology: C.S.L. Investigation: C.S.L., J.G., Y.L., M.X., J.H., B.V. Resources: D.C.H. and Y.Z. Formal analysis: C.S.L., J.G., M.L. Data Curation: C.S.L., J.G. Supervision: D.C.H. and Y.Z. Funding acquisition: D.C.H. and Y.Z. Writing – original draft preparation: C.S.L., J.G., D.C.H. and Y.Z. Writing – review and editing: C.S.L., J.G., D.C.H. and Y.Z.

## Methods

### List of antibodies

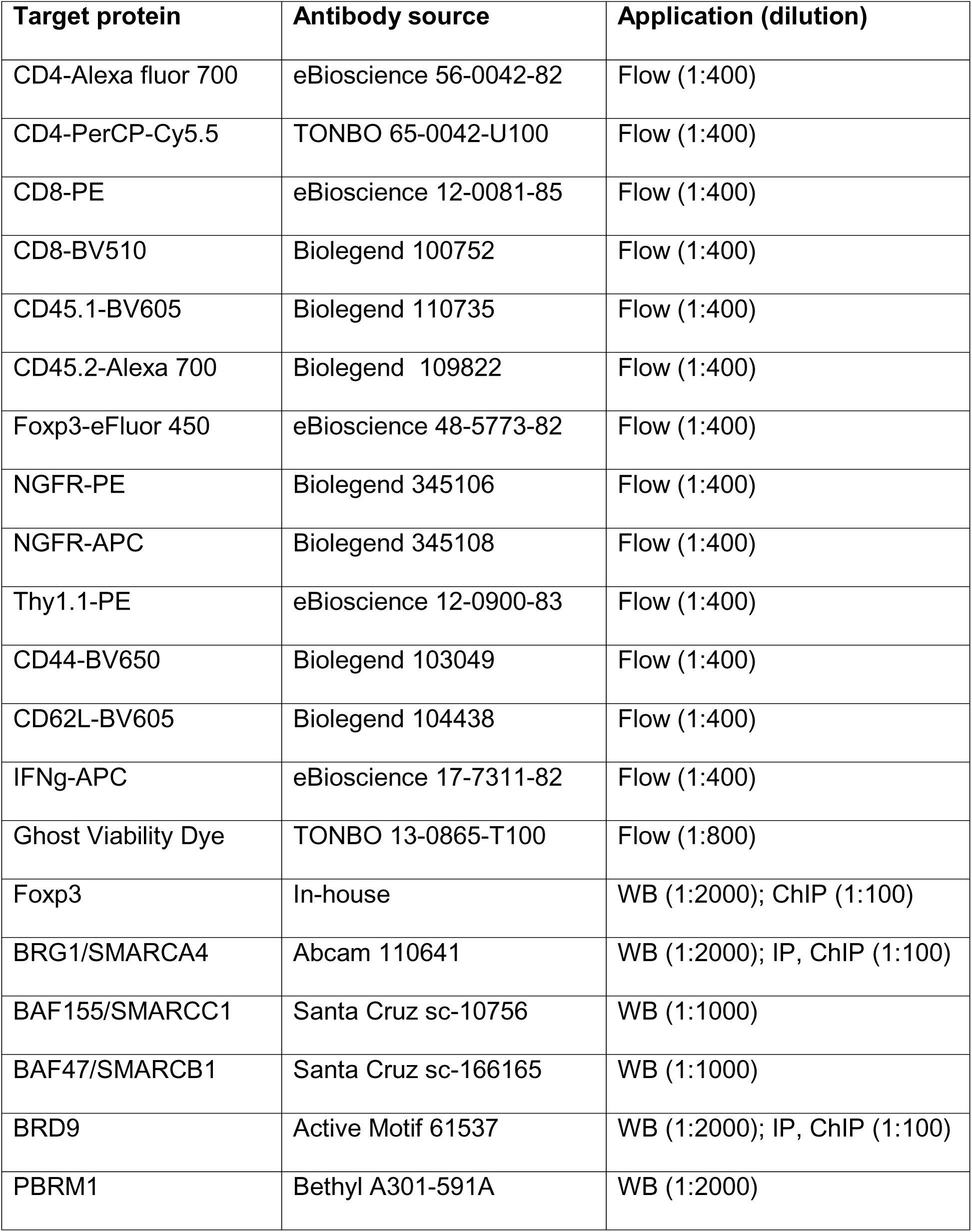
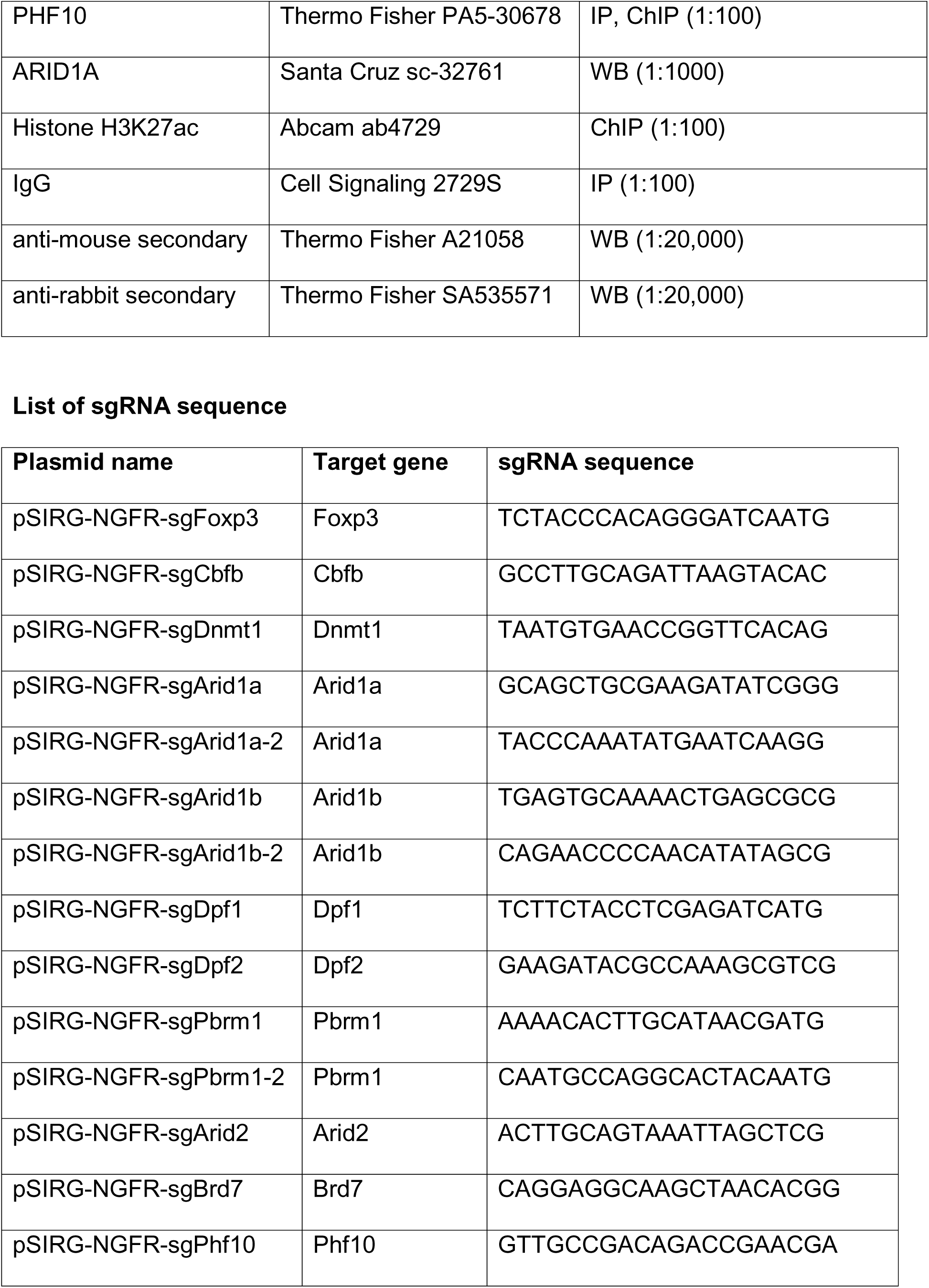
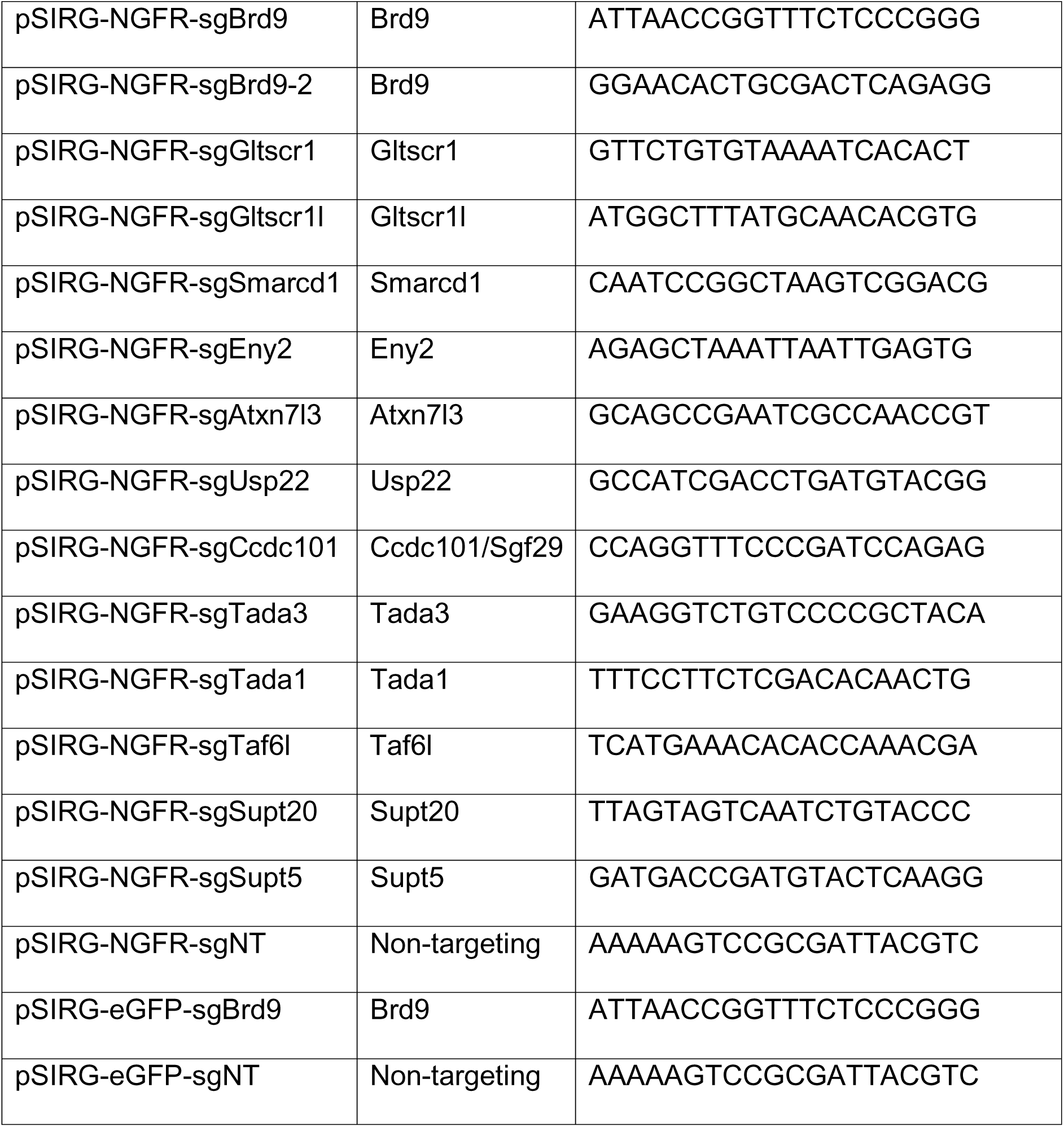

### Mice

C57BL/6 Rosa-Cas9/Foxp3^Thy1.1^ mice were generated by crossing Rosa26-LSL-Cas9 knockin mice(Platt et al., 2014) (The Jackson Laboratory #024857) with Foxp3^Thy1.1^ reporter mice(Liston et al., 2008). Male Cas9/Foxp3^Thy1.1^ mice at 8-12 weeks age were used to isolate Tregs for the CRISPR screen, and no gender preference was given for other experiments. C57BL.6 Ly5.1+ congenic mice and Rag1-/- mice purchased from the Jackson Laboratory were used for Treg suppression assay and adoptive T cell transfer in colitis and tumor models. All mice were bred and housed in the specific pathogen-free facilities at the Salk Institute for Biological Studies and were conducted under the regulation of the Institutional Animal Care and Use Committee (IACUC) and institutional guidelines.

### Retroviral vectors and sgRNA library construction

Self-inactivating retroviral vector pSIRG-NGFR was generated by modifying pSIR-dsRed-Express2(Fujita and Fujii, 2014) (Addgene #51135), which enables us to clone sgRNA as efficient as lentiCRISPRv2, to enrich transduced cells via magnetic beads isolation, and to perform intracellular staining without losing transduced reporter marker. We first mutated all BbsI sites in pSIR-dsRed-Express2, then inserted a sgRNA expressing cassette containing the U6 promoter, guide RNA scaffold and a 500bp filler embedded with BbsI cloning site. The dsRed cassette was replaced by cDNA sequence of human NGFR with truncated intracellular domain. We also generated pSIRG vector with eGFP (pSIRG-eGFP) for the purpose of T cells transfer in tumor study, minimizing potential immune rejection. The pSIRG-eGFP was generated by cutting pSIRG-NGFR with XcmI to remove NGFR cassette and replaced by eGFP cDNA by Gibson cloning. For cloning single guide RNA into the pSIRG vector, an annealed sgRNA oligos can be directly inserted into BbsI-digested pSIRG-NGFR by T4 ligation similar to the cloning method utilized by lentiCRISPRv2(Sanjana et al., 2014). To create a pooled sgRNA library in pSIRG-NGFR, we first amplified sgRNA sequences from an optimized mouse CRISPR knockout library lentiCRISPRv2-Brie (Addgene #73632). A total of eight 50 μL PCR reactions were performed to maximize coverage of sgRNA complexity. Each 50 μL PCR reaction contained Q5 High-Fidelity DNA polymerase and buffer (NEB #M0491), 15ng of lentiCRISPRv2-Brie, and targeted primers (Forward: GGCTTTATATATCTTGTGGAAAGGACGAAACACCG, Reverse: CTAGCCTTATTTTAACTTGCTATTTCTAGCTCTAAAAC). PCR was performed at 98°C denature, 67°C annealing, 72°C extension for 12 cycles. The sgRNA library amplicons were then combined and separated in 2 % agarose gel, and purified by the QIAquick Gel Extraction Kit (Qiagen #28704). The purified sgRNA amplicons was inserted into the BbsI-digested pSIRG-NGFR by NEBuilder HIFI assembly (NEB #E2621S). The sgRNA representative of the retroviral CRISPR library (pSIRG-NGFR-Brie) was validated by deep sequencing and comparing to the original lentiCRISPRvs-Brie. The coverage of the new pSIRG-NGFR sgRNA library was evaluated by the PinAPL-Py program (Spahn et al., 2017) (see Extended Data Figure 1).

### T cell isolation and culture

For large scale Treg culture, we first expanded Treg in Rosa-Cas9/Foxp3^Thy1.1^ mice by injecting IL-2:IL-2 antibody immune complex according protocol described in Webster KE et. al(Webster et al., 2009). Spleen and lymph node Tregs were labeled with PE-conjugated Thy1.1 antibody and isolated by magnetic selection using anti-PE microbeads (Mitenyl #130-048-801). All isolated Tregs were activated by plate bound anti-CD3 and anti-CD28 antibodies and cultured with X-VIVO 20 media (LONZA #04-448Q) supplemented by 1X Pen/Strep, 1X Sodium pyruvate, 1X HEPES, 1X GlutaMax, 55 μM beta-mercaptoethanol in the presence of IL-2 at 500 U/mL. For experiments with BRD9 degradation, Tregs were treated at day 0 with 2.5μM dBRD9 (Tocris #6606) and cultured for four days for RNA- and ChIP-seq and 0.16-10μM treated at day 0 and cultured dBRD9 for four days for Foxp3 MFI, cell viability and cell proliferation assays. Live cells were enriched by Ficoll-Paque 1.084 (GE Health 17-5446-02) for RNA-seq and ChIP-seq.

### Retroviral production and T cell transduction

HEK293T cells were seeded in 6-wells plate at 0.5 million cells per 2mL DMEM media supplemented by 10% FBS, 1% Pen/Strep, 1X GlutaMax, 1X Sodium Pyruvate, 1X HEPES, and 55 μM beta-mercaptoethanol. One day later, cells from each well was transfected with 1.2 μg of targeting vector pSIRG-NGFR and 0.8 μg of packaging vector pCL-Eco (Addgene, #12371) by using 4 μL of FuGENE HD transfection reagent (Promega #E2311) according manufactured protocol. Cell culture media was replaced by 3 mL fresh DMEM complete media at 24 hours and 48 hours after transfection. The retroviral supernatant was collected at 48 and 72 hours post transfection for T cell infection. For experiments with CRISPR sgRNA targeted knockdown, Cas9+ Tregs were first seeded in 24-wells plate coated with CD3 and CD28 antibodies. At 24 hour post-activation, 70% of Treg media from each well was replaced by retroviral supernatant, supplemented with 4 μg/mL Polybrene (Milipore # TR-1003-G), and spun in a benchtop centrifuge at 1,258 x g for 90 minutes at 32°C. After centrifugation, Treg media was replaced with fresh media supplemented with IL-2 and cultured for another three days. Transduced cells were analyzed for Foxp3 and cytokine expression in eBioscience Fix/Perm buffer (eBioscience #00-5523-00) using flow cytometry. Transduced NGFR+ cells were FACS-sorted for subsequent RNA- and ChIP-seq experiments.

### Genome-wide CRISPR screen in Treg

Approximately 360 million Treg cells isolated from Rosa-Cas9/Foxp3^Thy1.1^ mice were used for the Treg screen. On day 0, Tregs were seeded at 1×10^6^ cells/mL into 24-wells plate coated with anti-CD3/28 and cultured with X-VIVO complete media with IL-2 (500 U/ml). On day 1, sgRNA retroviral library transduction was performed with a MOI<0.2. On day 3, approximately 4 million (∼50X coverage) NGFR+ transduced cells were collected in three replicates as the starting state sgRNA input. Treg cells reached confluence on day 4. NGFR+ transduced cells were isolated via magnetic selection by anti-PE beads (Mitenyl #130-048-801), and then plated onto new 24-wells plates coated with anti-CD3/CD28, and cultured in X-VIVO complete media with IL-2 (500 U/ml). On day 6, approximately 4 million NGFR+ transduced cells were collected in three replicates as the ending state sgRNA output. The remaining cells were fixed, permeabilized, and stained for intracellular Foxp3. Approximately 2 million Foxp3^High^ (top 20%) and 2 million Foxp3^Low^ (bottom 20%) cell populations were sorted in three replicates by a FACS Aria cell sorter for genomic DNA extraction and library construction.

### Preparation of sgRNA amplicons for Next-Generation Sequencing

To extract genomic DNA, we first lysed cells with homemade digestion buffer (100mM NaCl, 10mM Tris, 25mM EDTA, 0.5% SDS, 0.1mg/mL Proteinase K) overnight in 50 °C. On the following day, the lysed sample was mixed with phenol: chloroform: isoamyl alcohol (25:24:1, v/v) in 1:1 ratio, and spun at 6000rpm for 15 min at room temperature. The supernatant containing genomic DNA was transferred into a new tube and mixed with twice volume of 100% ethanol, then spun at 12,500 rpm for 5 min in room temperature to precipitate DNA. Supernatant was removed, and the precipitated DNA was dissolved in ddH_2_O. DNA concentration was measured by Nanodrop. To generate sgRNA amplicons from extracted genomic DNA, we used a two-step PCR protocol which was adopted from the protocol published by *Shalem et. al. (Shalem et al., 2014)*. We performed eight 50 μL PCR reactions containing 2 μg genomic DNA, NEB Q5 polymerase, and buffer, and targeted primers (Forward: GGCTTTATATATCTTGTGGAAAGGACGAAACACCG, Reverse: CTAGCCTTATTTTAACTTGCTATTTCTAGCTCTAAAAC). PCR was performed at 98°C denature, 70°C annealing, 15s extension for 20 cycles. The products from the first PCR were pooled together, and purified by AMPure XP SPRI beads according to manufacturer’s protocol, and quantified by Qubit dsDNA HS assay. For the second round PCR, we performed eight 50 μL PCR reactions containing 2 ng purified 1^st^ round PCR product, barcoded primer (see primer set from *(Shalem et al., 2014)*, Priming site of reverse primer was changed to CTTCCCTCGACGAATTCCCAAC), NEB Q5 polymerase, and buffer. PCR was performed at 98°C denature, 70°C annealing, 15 s extension for 12 cycles. The 2^nd^ round PCR products were pooled, purified by AMPure XP SPRI beads, quantified by Qubit dsDNA HS assay, and sequenced by NEXTSeq sequencer at single end 75 bp (SE75).

### Data analysis of pooled CRISPR screen

The screening hit identification and quality control was performed by MAGeCK-VISPR program(Li et al., 2015; Li et al., 2014a). The abundance of sgRNA from a sample fastq file was first quantified by MAGeCK “Count” module to generate a read count table. For hit calling, we used MAGeCK “test” module to generate a gene-ranking table that reporting RRA gene ranking score, p-value, and log2 fold change. The size factor for normalization was adjusted according to1000 non-targeting control assigned in the screen library. All sgRNAs that are zero read were removed from RRA analysis. The log2 fold change of a gene was calculated from a mean of 4 sgRNA targeting per gene. The scatter plots showing the screen results were generated by using the R script EnhancedVolcano (https://github.com/kevinblighe/EnhancedVolcano). The R script that generated the sgRNA distribution histogram was provided by E. Shifrut and A. Marson (UCSF)(Shifrut et al., 2018). A gene list from Foxp3 regulators (either positive or negative) without affecting cell proliferation was subjected to Gene Ontology analysis using Metascape(Zhou et al., 2019). Genes were analyzed for enrichment for Functional Set, Pathway, and Structural Complex.

### In vitro Treg suppression assay

Tregs were transduced by retrovirus expressing sgRNA targeting gene of interest and cultured in X-VIVO complete media supplemented with IL-2 (500 U/ml). Four days after transduction, transduced cells were sorted and mixed with FACS sorted CD45.1+ naive CD4 T cells (CD4^+^ CD25^−^ CD44^Low^ CD62L^High^) labeled with CellTrace Violet (Thermo Fisher Scientific #C34571) in different ratio in the presence of irradiated T cell depleted spleen cells as antigen-presenting cells (APC). Three days later, Treg suppression function was measured by the percentage of non-dividing cells within the CD45.1^+^ effector T cell population. For dBRD9 treatment experiment, dBRD9 was first dissolved in DMSO (10 mM stock) and added into Treg:Teff:APC mixture at 2.5μM. For Foxp3 overexpression rescue experiment, Tregs were first transduced with sgNT or sgBrd9 at 24 hour post-activation, and then transduced with MIGR empty vector or MIGR-Foxp3 at 48 hour post-activation. Double transduced Tregs were FACS sorted on day 4 based on NGFR+ and GFP+ markers and then mixed with CellTrace labeled effector T cells in the presence of APC. Treg suppression readout was measured after three days of co-culture.

### Adoptive T cells transfer-induced colitis model

Tregs were transduced by retrovirus expressing sgRNA targeting gene of interest, and cultured in X-VIVO complete media and IL-2 (500 U/ml). Four days after transduction, the NGFR+ transduced Treg cells were FACS sorted before transferred into recipient mice. To induce colitis, 2 million effector T cells (CD45.1^+^ CD4^+^ CD25^−^ CD45RB^High^) and 1 million sgRNA knockout Tregs (CD45.2^+^ CD4^+^ Thy1.1^+^ NGFR^+^) were mixed together and transferred into Rag1 knockout recipient mice. The body weight of recipient mice was monitored weekly for signs of wasting symptoms. Mice were harvested 7 weeks after T cell transfer. Spleens were used for profiling immune cell populations by FACS. Colons were collected for histopathological analysis.

### Colon histopathological analysis

Histopathological analysis was performed in a blinded manner and scored using the following criteria. Eight parameters were used that include (i) the degree of inflammatory infiltrate in the LP (0-3); (ii) Goblet cell loss (0–2); (iii) reactive epithelial hyperplasia/atypia with nuclear changes (0–3); (iv) the number of IELs in the epithelial crypts (0–3); (v) abnormal crypt architecture (distortion, branching, atrophy, crypt loss) (0–3); (vi) number of crypt abscesses (0–2); (vii) mucosal erosion to frank ulcerations (0–2) and (viii) submucosal spread to transmural involvement (0-2). The severity of lesion was scored independently in 3 regions (proximal, middle and distal colon) over a maximal score of 20. The overall colitis score was based as the average of each regional score (maximal score of 20).

### Adoptive T cells transfer and MC38 tumor model

Similar to the “Adoptive T cells transfer-induced colitis model”, Tregs were activated in vitro and transduced with pSIRG-eGFP expressing sgNT or sgBrd9. Four days after transduction, the eGFP+ transduced Treg were FACS sorted. Concurrently, Treg depleted CD4 and CD8 T cells isolated from Rosa-Cas9/Foxp3^Thy1.1^ mice were used as effector T cells. A total of 1 million pSIRG-sgRNA transduced eGFP+ Tregs, 1 million effector CD8 T cells, and 2 million Treg-depleted CD4 T cells were mixed and transferred into Rag1 knockout recipient mice. on the following day, mice were implanted with 0.5 million MC38 cells (a gift from the laboratory of Dr. Susan Kaech) by subcutaneous injection on the flank of mouse. When palpable tumor appeared, tumor size was measured every two day by electronic calipers. At the end point, spleen and tumor were collected for immune profiling. For tumor processing, tumor tissues were minced into small pieces and digested with 0.5 mg/mL Collagenase IV (Sigma #C5138) and DNAase I (Roche #4716728001) for 20 minutes and passed through 0.75 μm cell strainer to collect single cell suspension. Isolated cells were stimulated with PMA/Ionomycin and Golgi plug for 5 hours, and then were subjected to Foxp3 and cytokines staining with eBioscience Fix/Perm buffer (eBioscience #00-5523-00).

### Nuclear protein extraction

Nuclear lysates were collected from Treg cells following a revised Dignam protocol(Andrews and Faller, 1991). After cellular swelling in Buffer A (10 mM Hepes pH 7.9, 1.5 mM MgCl_2_, 10 mM KCl) supplemented with 1 mM DTT, 1 mM PMSF, 1 µM pepstatin, 10 µM leupeptin and 10 µM chymostatin, cells were lysed by homogenization using a 21-gauge needle with six to eight strokes. If lysis remained incomplete, cells were treated with 0.025 - 0.05% Igepal-630 for ten minutes on ice prior to nuclei collection. Nuclei were spun down at 700 x g for five minutes then resuspended in Buffer C (20 mM Hepes pH 7.9, 20% glycerol, 420 mM NaCl, 1.5 mM MgCl_2_, 0.2 mM EDTA) supplemented with 1 mM DTT, 1 mM PMSF, 1 µM pepstatin, 10 µM leupeptin and 10 µM chymostatin. After thirty minutes of end-to-end rotation at 4°C, the sample was clarified at 21,100 x g for ten minutes. Supernatant was collected, flash frozen in liquid nitrogen and stored in the −80°C freezer.

### Co-Immunoprecipitation

Nuclear lysates were thawed on ice then diluted with two-thirds of original volume of 50 mM Tris-HCl pH 8, 0.3% NP-40, EDTA, MgCl_2_ to bring down the NaCl concentration. Proteins were quantified using Biorad DC Protein Assay (Cat #5000112) according to manufacturer’s instructions. For the co-IP reaction, 200-300 µg of proteins were incubated with antibody against normal IgG, SMARCA4, BRD9, ARID1A or PHF10 overnight at 4°C, with end-to-end rotation. Precipitated proteins were bound to 50:50 Protein A: Protein G Dynabeads (Invitrogen) for one to two hours and washed extensively with IP wash buffer (50 mM Tris pH 8, 150 mM NaCl, 1 mM EDTA, 10% glycerol, 0.5% Triton X100). Proteins were eluted in SDS-PAGE loading solution with boiling for five minutes and analyzed by western blotting.

### Western blot

Protein samples were run on 4-12% Bis-Tris gels (Life Technologies). After primary antibody incubation which is typically done overnight at 4°C, blots were probed with 1:20,000 dilution of fluorescently-labeled secondary antibodies in 2% BSA in PBST (1X Phospho-buffered saline with 0.1% Tween-20) for an hour at room temperature (RT). Fluorescent images were developed using Odyssey and analyzed using Image Studio 2. Protein quantitation was performed by first normalizing the measured fluorescence values of the proteins of interest against the loading control (TBP) then normalizing against the control sample (vehicle treated).

### RNA-seq sample preparation

RNA from 1-3 x 10^6^ cells was extracted and purified with TRIzol reagent (Thermo Fisher) according to manufacturer’s instructions. RNA-seq libraries were prepared using Illumina TruSeq Stranded mRNA kit following manufacturer’s instructions with 5 µg of input RNA.

### RNA-seq analysis

Single-end 50 bp reads were aligned to the mouse genome mm10 using STAR alignment tool (V2.5)(Dobin et al., 2013). RNA expression was quantified as raw integer counts using analyzeRepeats.pl in HOMER(Heinz et al., 2010) using the following parameters: -strand both -count exons -condenseGenes -noadj. To identify differentially expressed genes, we performed getDiffExpression.pl in HOMER, which uses the DESeq2 R package to calculate the biological variation within replicates. Cut-offs were set at log2 FC = 0.585 and FDR at 0.05 (Benjamin-Hochberg). Principal Component Analysis (PCA) was performed with the mean of transcript per million (TPM) values using Cluster 3.0 with the following filter parameters: at least one observation with absolute value equal or greater than two and gene vector of four. TPM values were log transformed then centered on the mean.

### Gene Set Enrichment Analysis

GSEA software(Mootha et al., 2003; Subramanian et al., 2005) was used to perform the analyses with the following parameters: number of permutations = 1000; enrichment statistic = weighted; and metric for ranking of genes = difference of classes (Input RNA-seq data was log-transformed). For Figure 5G, input RNA-seq data contained the normalized log-transformed reads of the 1,325 differentially expressed genes (DEGs) in sgFoxp3/sgNT Tregs. The compiled gene list included GSEA Gene Ontology, Immunological Signature, Curated Gene, and the up- and down-regulated DEGs in sgBrd9/sgNT Tregs. The resulting normalized enrichment scores and FWER p values were combined to generate the graph.

### ChIP-seq sample preparation

Treg cells were collected and cross-linked first in 3 mM disuccinimidyl glutarate (DSG) in 1X PBS for thirty minutes then in 1% formaldehyde for another ten minutes, both at RT, for chromatin binding protein ChIP or in 1% formaldehyde only for histone modification ChIP. After quenching the excess cross-linker with a final concentration of 125 mM glycine, the cells were washed in 1X PBS, pelleted, flash-frozen in liquid nitrogen, and stored at −80°C. Cell pellets were thawed on ice and incubated in lysis solution (50 mM HEPES-KOH pH 8, 140 mM NaCl, 1 mM EDTA, 10% glycerol, 0.5% NP40, 0.25% Triton X-100) for ten minutes. The isolated nuclei were washed with wash solution (10 mM Tris-HCl pH 8, 1 mM EDTA, 0.5 mM EGTA, 200 mM NaCl) and shearing buffer (0.1% SDS, 1 mM EDTA, 10 mM Tris-HCl pH 8) then sheared in a Covaris E229 sonicator for ten minutes to generate DNA fragments between ∼ 200-1000 base pairs (bp). After clarification of insoluble material by centrifugation, the chromatin was immunoprecipitated overnight at 4°C with antibodies against Foxp3, SMARCA4, BRD9, PHF10 or H3K27ac. The next day, the antibody bound DNA was incubated with Protein A+G Dynabeads (Invitrogen) in ChIP buffer (50 mM HEPES-KOH pH 7.5, 300 mM NaCl, 1 mM EDTA, 1% Triton X-100, 0.1% DOC, 0.1% SDS), washed and treated with Proteinase K and RNase A. Cross-linking was reversed by incubation at 55°C for two and a half hours. Purified ChIP DNA was used for library generation (NuGen Ovation Ultralow Library System V2) according to manufacturer’s instructions for subsequent sequencing.

### ChIP-seq analysis

Single-end 50 bp or paired-end 42 bp reads were aligned to mouse genome mm10 using STAR alignment tool (V2.5)(Dobin et al., 2013). ChIP-Seq peaks were called using findPeaks within HOMER using parameters for histone (-style histone) or transcription factor (-style factor) (Christopher Benner, HOMER, http://homer.ucsd.edu/homer/index.html, 2018). Peaks were called when enriched > two-fold over input and > four-fold over local tag counts, with FDR 0.001 (Benjamin-Hochberg). For histone ChIP, peaks within a 1000 bp range were stitched together to form regions. ChIP-Seq peaks or regions were annotated by mapping to the nearest TSS using the annotatePeaks.pl command. Differential ChIP peaks were found by merging peaks from control and experiment groups and called using getDiffExpression.pl with fold change ≥1.5 or ≤-1.5, Poisson p value < 0.0001.

### Motif analysis

Sequences within 200 bp of peak centers were compared to motifs in the HOMER database using the findMotifsGenome.pl command using default fragment size and motif length parameters. Random GC content-matched genomic regions were used as background. Enriched motifs are statistically significant motifs in input over background by a p-value of less than 0.05. P-values were calculated using cumulative binomial distribution.

### ATAC-seq sample preparation

ATAC-seq was performed according to previously published protocol(Buenrostro et al., 2013). Briefly, 50,000 Treg cells were collected in duplicates and washed first with cold 1X PBS then with Resuspension buffer (RSB; 10 mM Tris-HCl pH 7.4, 10 mM NaCl, 3 mM MgCl_2_). Cells were lysed in RSB supplemented with 0.1% Igepal-630 and nuclei were isolated by centrifugation at 500 x g for ten minutes. Nuclei were incubated with Tn5 transposase in Tagment Buffer (Illumina) for thirty minutes at 37°C. Purified DNA was ligated with adapters, amplified and size selected using AMPure XP beads (Beckman) for sequencing. Library DNA was sequenced using paired end 42 bp reads.

### ATAC-seq analysis

Paired end 42 bp reads were aligned to mouse genome mm10 using STAR alignment tool (V2.5). ATAC-seq peaks were called using findPeaks within HOMER using the style parameter dnase. Peaks were called when enriched > four-fold over genomic background and > four-fold over local tag counts, with FDR 0.001 (Benjamin-Hochberg).

