## Supplementary Materials for "A genome-wide CRISPR screen reveals a role for the BRD9-containing non-canonical BAF complex in regulatory T cells"

Loo et al Figure S1

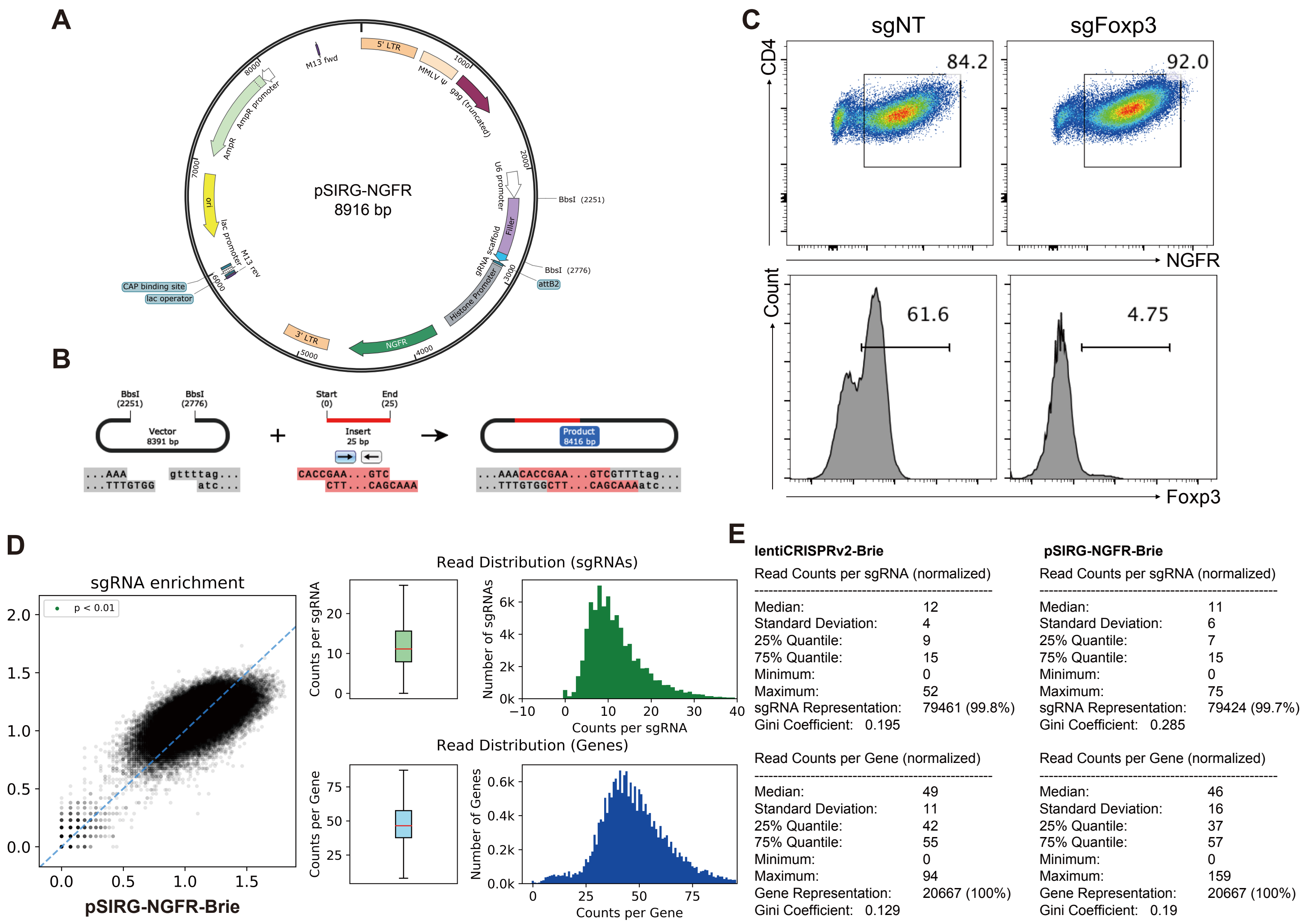


**Figure S1. Construction of a retroviral sgRNA CRISPR library (pSIRG-NGFR-Brie), Related to Figure 1.**

**A,** The map of pSIRG-NGFR. A self-inactivating retroviral vector containing a sgRNA expressing cassette and a truncated human NGFR surface marker. **B,** Overview of the process to clone a sgRNA into pSIRG-NGFR. A pair of annealed sgRNA oligomers can be directly cloned into BbsI-digested pSIRG-NGFR by T4 ligation. **C,** Validation of the transduction and knockout efficiency of pSIRG-NGFR. Cas9-expressing naïve CD4 T cells were transduced with either non-targeting control virus (sgNT) or Foxp3 targeting virus (sgFoxp3) in the presence of TGF-β and IL-2 for Foxp3 induction. NGFR and Foxp3 expression were measured by FACS 3 days post-infection. **D,** Correlation of sgRNA representation comparing lentiCRISPRv2-Brie library to pSIRG-NGFR-Brie library (left). Read distribution of sgRNAs and genes in pSIRG-NGFR-Brie (right). **E,** Statistics of sgRNAs and genes represented in lentiCRISPRv2-Brie and pSIRG-NGFR-Brie. Quantification of sgRNAs and genes was computed by PinAPL-Py program.

### Loo, et al. Figure S2

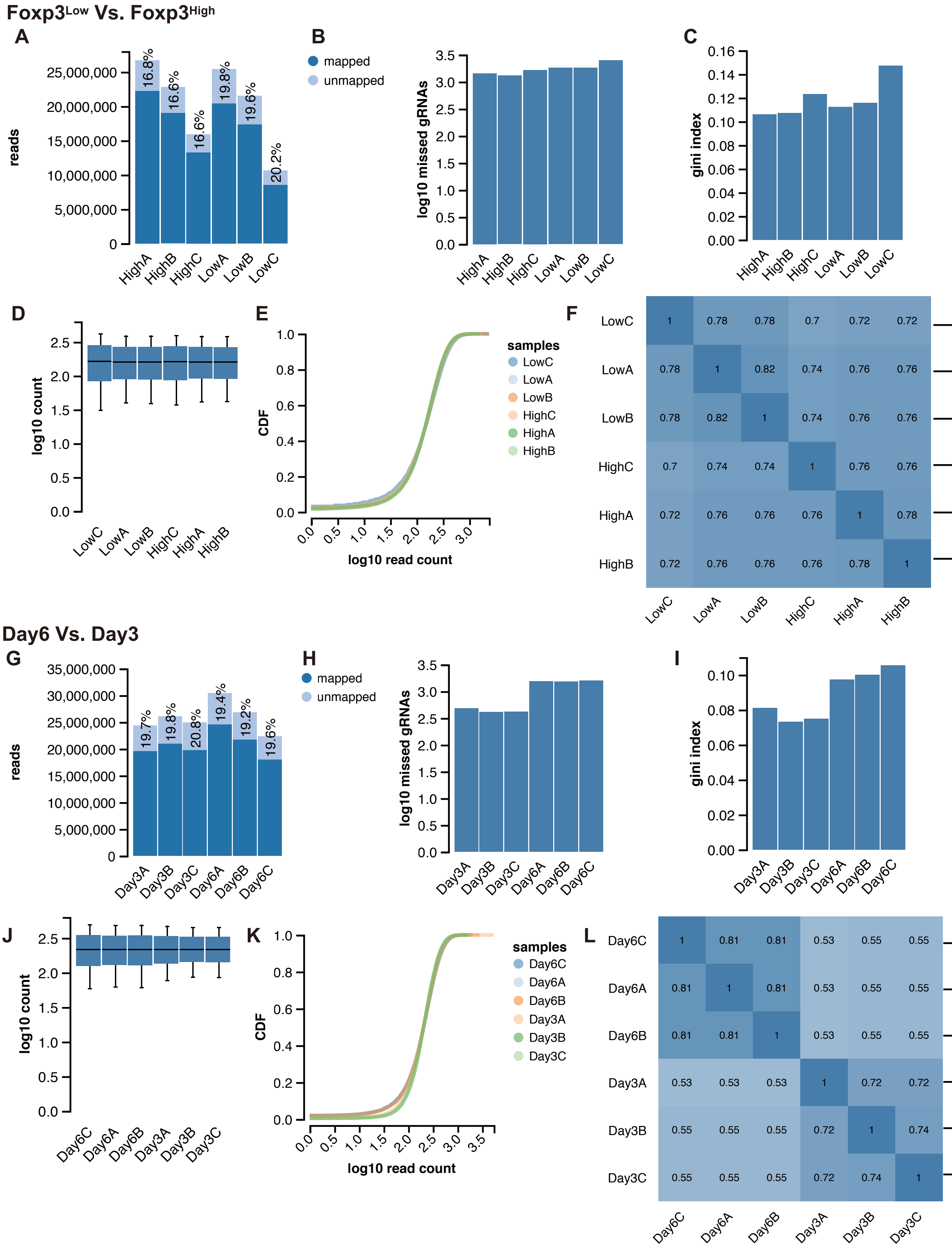

**Figure S2. Quality control analysis of samples generated from the Treg screen, Related to Figure 1.**

Quality control analysis of samples comparing between Foxp3Low and Foxp3High populations (**A-F**) or between Day 6 and Day 3 NGFR+ transduced populations (**G-L**). **A**, **G**, Mapped (dark blue) and unmapped (light blue) reads for each sample. Percentage of unmapped reads is labeled on each bar. **B**, **H**, Number of missed gRNAs with zero mapped reads. **C**, **I**, Gini Index for each sample measuring inequality between read counts. **D**, **J**, Distribution of normalized read counts for each sample. **E**, **K**, Cumulative distribution function of normalized read counts for each sample. **F**, **L**, Correlation between normalized log10 read counts of samples.

**Loo, et al. Figure S3**

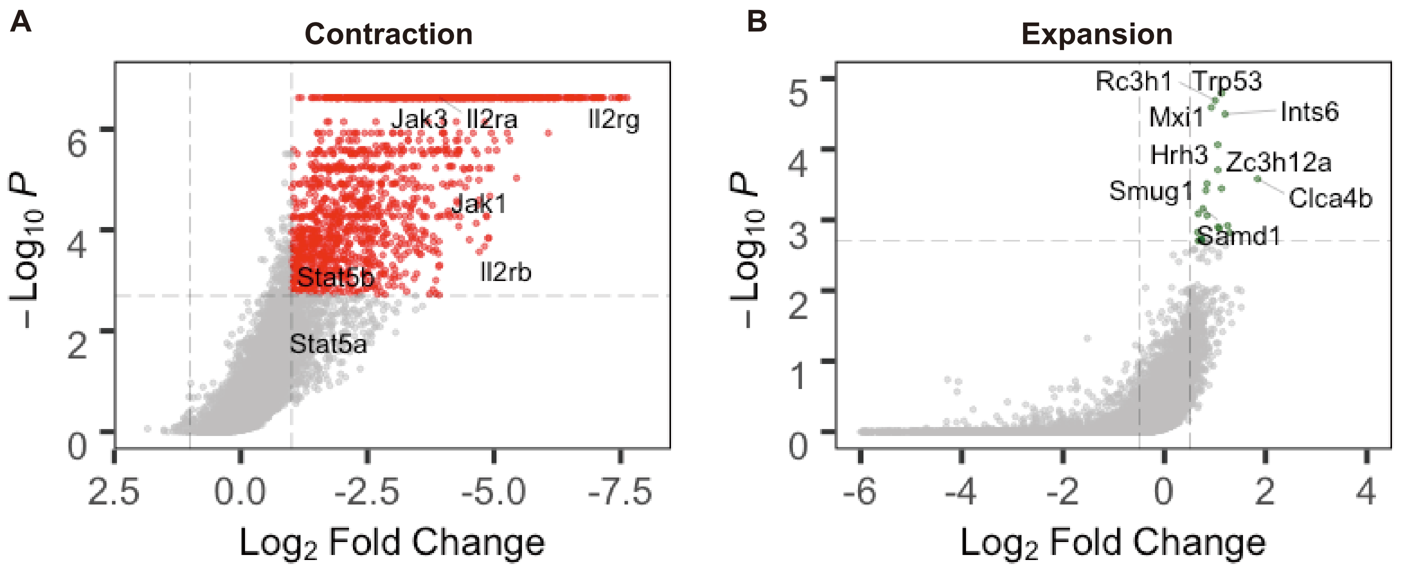


**Figure S3. Identification of genes that are regulating cell proliferation and survival in the Treg screen, Related to Figure 2.**

**A,B** Scatter plots showing genes enriched in the cell contraction pool (**A**) or cell expansion pool (**B**) by comparing NGFR+ transduced cells on day 6 to NGFR+ transduced cells on day 3 during the Treg screen. Cutoff was set for contraction is P-value <0.002 and LFC>1 (Red dots), whereas cutoff for expansion was set P value <0.002 and LFC >0.5 (Green dots).

Loo, et al. Figure S4

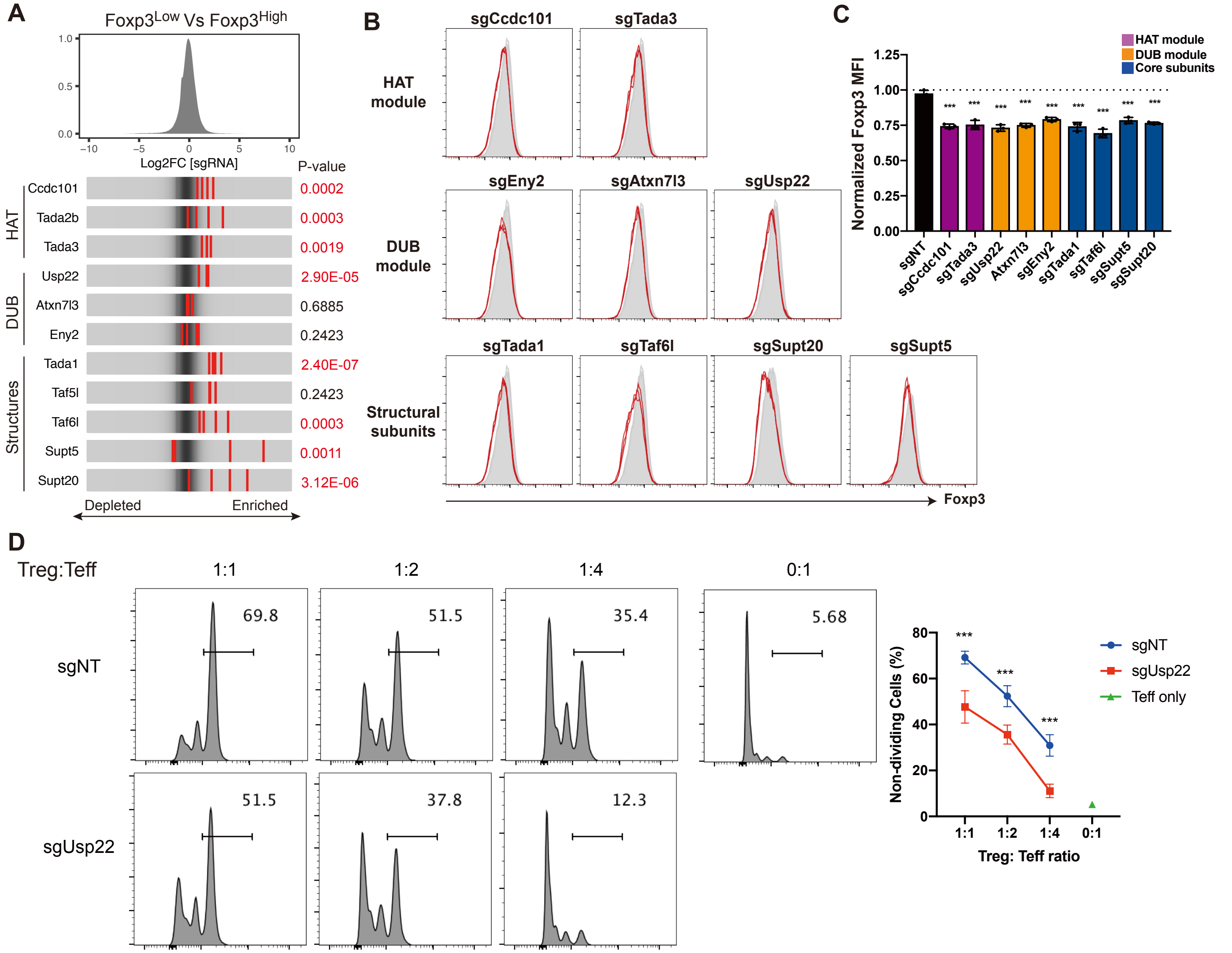


**Figure S4. The SAGA complex regulates Foxp3 expression and Treg suppressor activity, Related to Figure 2.**
**A**, Distribution of sgRNA Log2FC comparing Foxp3Low to Foxp3High. Red stripes represent sgRNAs from positive Foxp3 regulators. Genes with a P-value of less than 0.01 were shown in red. **B**, FACS plot of Foxp3 expression in Tregs transduced with sgRNAs against Ccdc101, Tada3, (HAT module), Eny2, Atxn7l3 and Usp22 (DUB module), and Tada1, Taf6l, Supt20, Supt5 (structural subunits) of SAGA complex (n=3 per group.). **C**, Mean fluorescent intensity (MFI) of Foxp3 in Tregs transduced with sgRNAs against SAGA subunits. **D**, In vitro suppression assay of Tregs transduced with sgUsp22. sgNT is non-targeting control. n=3 per group. Data represent mean ± s.d. Statistical analyses were performed using unpaired two-tailed Student's t test (***p<0.001).

**Loo, et al. Figure S5**

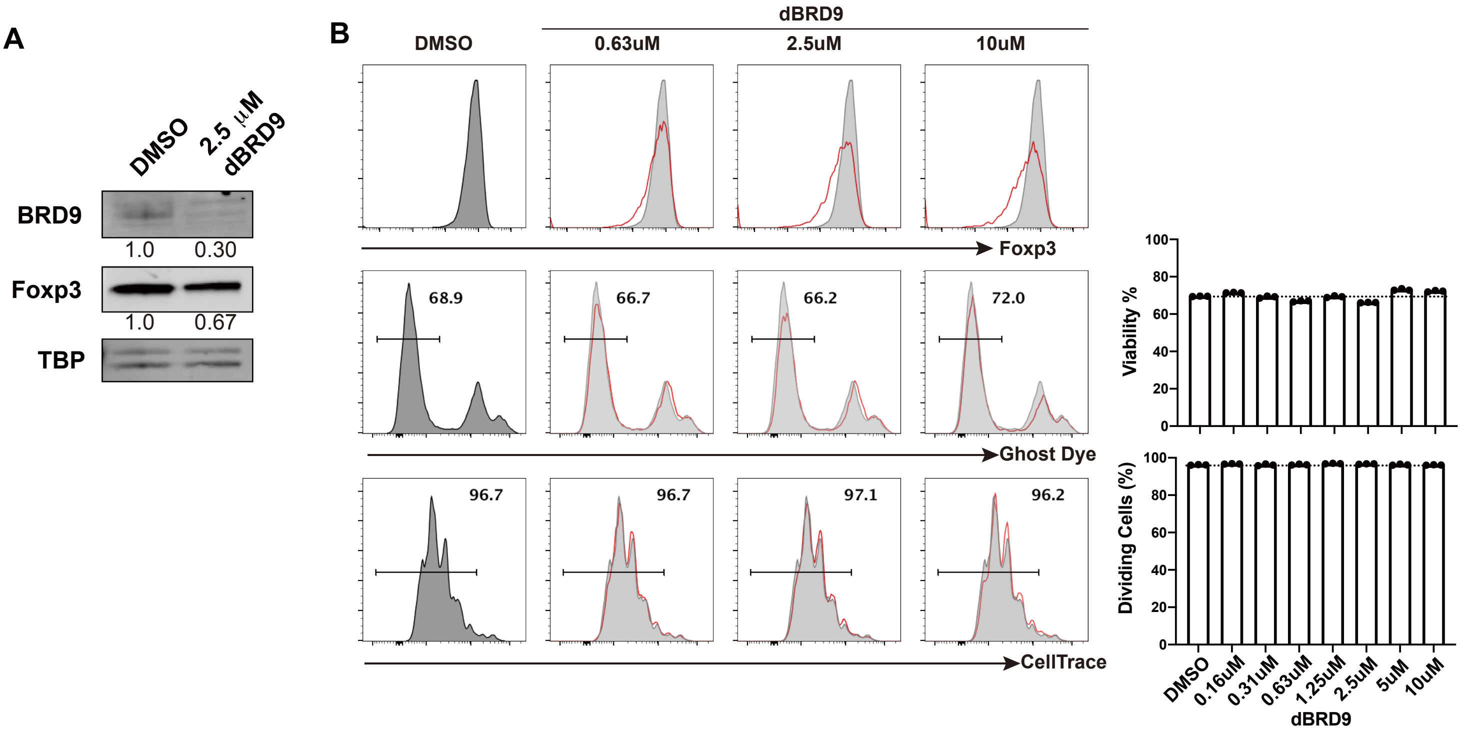


**Figure S5.** **BRD9 degrader dBRD9 reduces Foxp3 expression without affecting cell viability and proliferation, Related to Figure 3**

**A**, Immunoblotting analysis of BRD9, Foxp3, and TATA-binding protein (TBP) in nuclear lysates from Tregs treated with either DMSO or 2.5 μM dBRD9 for four days. Normalized protein levels are indicated. **B,** Foxp3 expression, cell viability labeled by Ghost Dye, and cell division determined by CellTrace dilution in Tregs after treatment of dBRD9 in increasing concentrations for 4 days (n=3 per group). Grey shade: DMSO. Red line: dBRD9. See also Figure 3E. Data represents mean ± sd. Statistical analyses were performed using unpaired two-tailed Student's t-test. (*$p<0.05$, **$p<0.01$, ***$p<0.001$).

**Loo, et al. Figure S6**

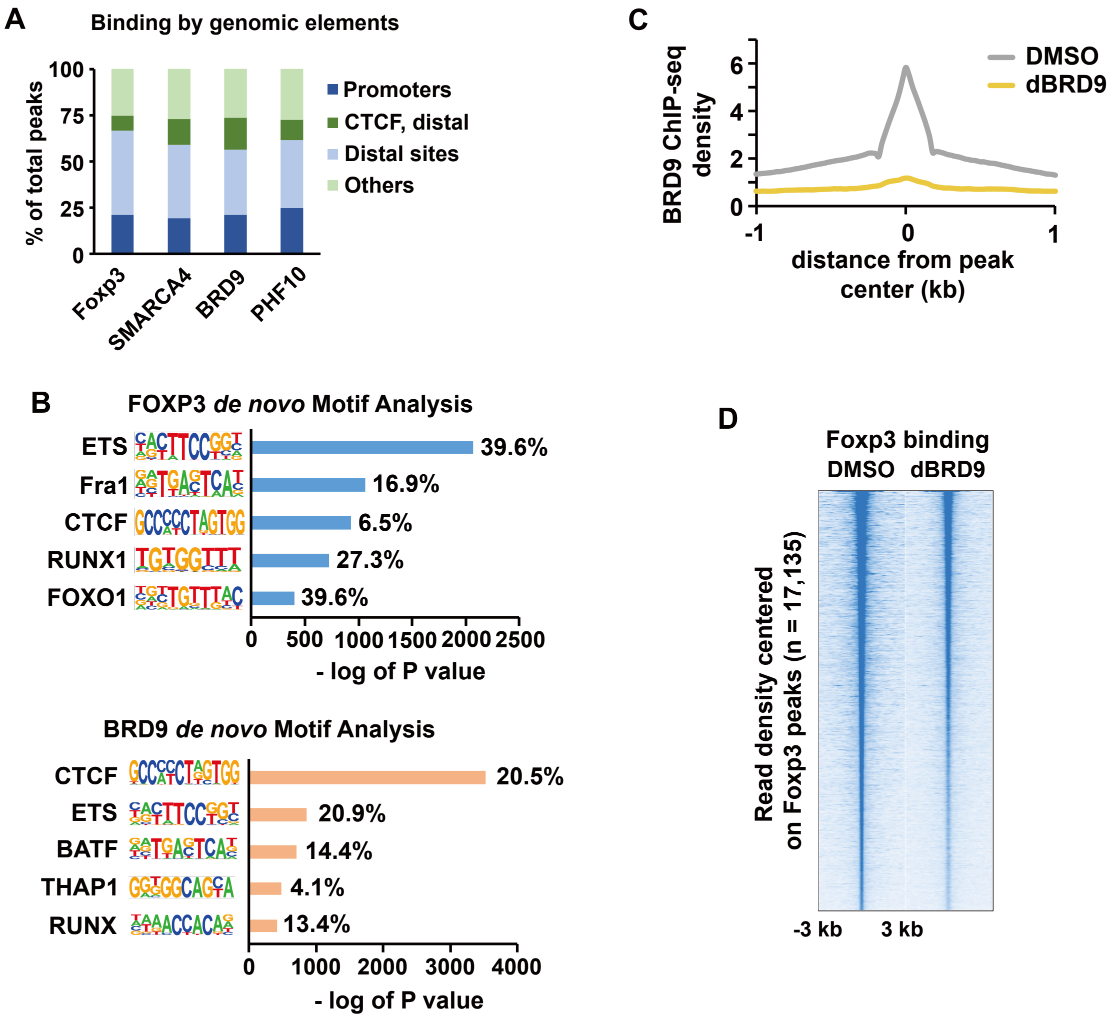


**Figure S6. BRD9 and Foxp3 co-localize on chromatin; BRD9 regulates Foxp3 binding to a subset of Foxp3 binding sites , Related to Figure 4.**

**A,** Stacked bar graph of sites bound by Foxp3, SMARCA4, BRD9, and PHF10 that localize to the indicated genomic elements. **B**, Bar graph showing the top five de novo motifs enriched at Foxp3 (top) and BRD9 (bottom) ChIP-seq peaks, the percentage of sites that contain the motif, and the negative log of P value (Binomial distribution against random genomic background). **C**, Histogram of BRD9 ChIP read density ± 1 kb surrounding the site peak center in DMSO and 2.5 μM dBRD9 treated nTregs. **D**, Heatmap of Foxp3 ChIP-seq signal in DMSO and dBRD9-treated nTregs ± 3 kb centered on Foxp3-bound sites in DMSO, ranked according to read density.

**Loo, et al. Figure S7**

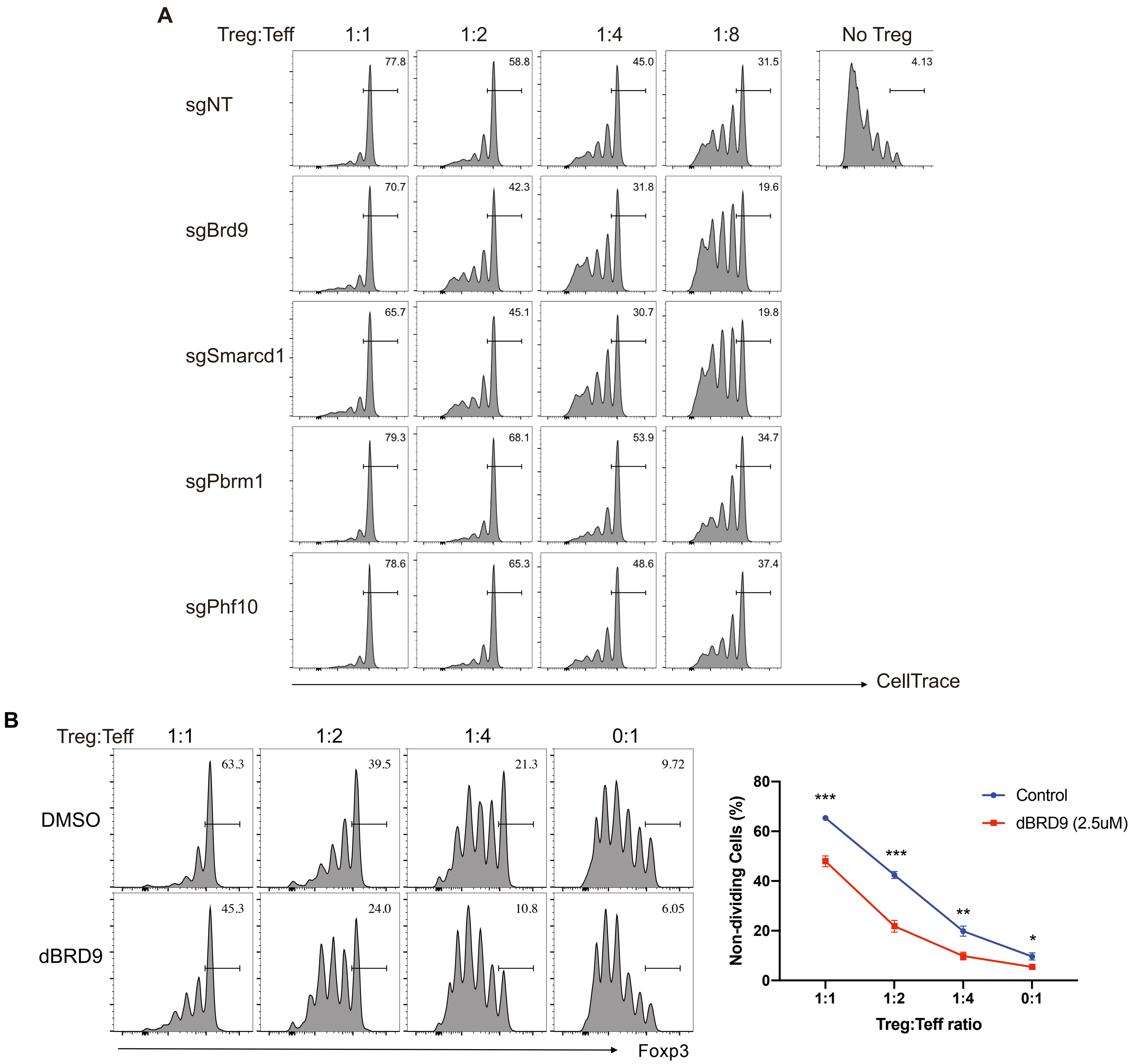


**Figure S7. sgRNA knockout or chemical degradation of GBAF or PBAF subunits alters Treg suppressor function, Related to Figure 6.**

**A,** In vitro suppression assay of Tregs with sgRNA knockout of Brd9, Smarcd1, Pbrm1, and Phf10 (n=3 per group, data represent mean ± s.d.). sgNT was used as non-targeting control. Also see Figure 6A. **B,** In vitro suppression assay using Tregs treated with dBRD9 or vehicle DMSO. Representative histograms of effector T cell divisions in different Treg:Teff ratios. (n=3 per group, data represent mean ± s.d.). Statistical analyses were performed using unpaired two-tailed Student's t test (ns: p>=0.05, *p<0.05, **p<0.01, ***p<0.001).

**Supplementary Tables**

**Table S1. List of genes in comparing Foxp3 Low to Foxp3 High populations identified in the CRISPR-Cas9 screen of Treg cells, Related to Figure 2.**

This excel file consists of four tabs. A complete gene list with statistical analysis comparing Foxp3 low and high population is shown in the first tab labeled “Foxp3 LowVsHigh”. Genes that met the cutoff criteria are shown in the second tab labeled “Foxp3 Pos Regulator” and in the third tab labeled “Foxp3 Neg Regulator”, representing Foxp3 positive and negative regulators, respectively. In the fourth tab, a complete list of raw read counts is available for each sgRNA for all three replicates of the screen.

**Table S2. List of genes List of genes comparing Day6 cells to Day3 cells for identifying genes involved in cell proliferation and survival, Related to Figure 2.**

A complete gene list with statistical analysis comparing cells on day 6 and day 3 is shown in the first tab labeled “Day6 Vs Day3”. Genes that met cutoff criteria are shown in the second tab labeled “Contraction” and in the third tab labeled “Expansion”, representing genes whose deletion led to cell population contraction or expansion, respectively. In the fourth tab, a complete list of raw read counts is available for each sgRNA for all three replicates of the screen.

**Table S3. List of genes in different categories identified in the CRISPR-Cas9 screen, Related to Figure 2.**

A summary of unique or shared genes identified in the Treg screen that regulate Foxp3 expression and/or cell contraction/expansion.

**Table S4. Gene ontology analysis of positive and negative Foxp3 regulators identified in the CRISPR-Cas9 screen of Treg cells, Related to Figure 2.**

Gene Ontology analysis of positive and negative Foxp3 regulators that do not affect Treg cell survival and proliferation (in tab 1 and tab 2, respectively) identified in the Treg screen was performed using Metascape.

**Table S5. GSEA enrichment analysis of Foxp3-dependent genes in Tregs, Related to Figure 5.**

List of the Gene Ontology, C2, Immunology, and BRD9-dependent gene lists that were probed against the RNA-seq expression data of Foxp3-dependent genes in sgFoxp3 and sgNT transduced Tregs.
